# Molecular signature of postmortem lung tissue from COVID-19 patients suggests distinct trajectories driving mortality

**DOI:** 10.1101/2021.11.08.467705

**Authors:** Anshul Budhraja, Anubhav Basu, Atish Gheware, Dasari Abhilash, Seesandra Rajagopala, Suman Pakala, Madhuresh Sumit, Animesh Ray, S Arulselvi, Purva Mathur, Aruna Nambirajan, Sachin Kumar, Ritu Gupta, Naveet Wig, Anjan Trikha, Randeep Guleria, Chitra Sarkar, Ishaan Gupta, Deepali Jain

## Abstract

The precise molecular mechanisms behind life-threatening lung abnormalities during severe SARS-CoV-2 infections are still unclear. To address this challenge, we performed whole transcriptome sequencing of lung autopsies from 31 patients suffering from severe COVID-19 related complications and 10 uninfected controls. Using a metatranscriptome analysis of lung tissue samples we identified the existence of two distinct molecular signatures of lethal COVID-19. The dominant “classical” signature (n=23) showed upregulation of unfolded protein response, steroid biosynthesis and complement activation supported by massive metabolic reprogramming leading to characteristic lung damage. The rarer signature (n=8) potentially representing “Cytokine Release Syndrome” (CRS) showed upregulation of cytokines such IL1 and CCL19 but absence of complement activation and muted inflammation. Further, dissecting expression of individual genes within enriched pathways for patient signature suggests heterogeneity in host response to the primary infection. We found that the majority of patients cleared the SARS-CoV-2 infection, but all suffered from acute dysbiosis with characteristic enrichment of opportunistic pathogens such as *Staphylococcus cohnii* in “classical” patients and *Pasteurella multocida* in CRS patients. Our results suggest two distinct models of lung pathology in severe COVID-19 patients that can be identified through the status of the complement activation, presence of specific cytokines and characteristic microbiome. This information can be used to design personalized therapy to treat COVID-19 related complications corresponding to patient signature such as using the identified drug molecules or mitigating specific secondary infections.

## Introduction

Despite numerous interventions, the novel coronavirus (SARS-CoV-2) continues to cause significant morbidity and mortality throughout the world. As of mid-August 2021, India alone has diagnosed over 32.2 million people with this virus, with over 400,000 fatalities (Dong et al., 2020). Though COVID-19 is believed to progress often asymptomatically or with only mild to moderate symptoms, primarily fever and dry cough, in many instances it can exacerbate acute pneumonia, especially in susceptible patients such as older individuals with metabolic, cardiovascular, and/or pulmonary comorbidities (Mehrian-Shai, 2020; Tay et al., 2020).

As it has been reported, SARS-CoV-2 enters the host cell using Angiotensin-Converting Enzyme-2 (ACE2) receptor, which binds to the viral spike (S) protein’s receptor binding domain (RBD) (Hoffmann et al., 2020). The viral genome is released into the cytoplasm once the viral envelope fuses with the host cell membrane in a Toll-like receptor-7 (TLR-7) dependent manner (Ahmadpoor & Rostaing, 2020). The virus uses its own RNA dependent RNA polymerase enzyme to replicate its genome (Sexton et al., 2016; Simmons et al., 2013). The replication-transcription complex (RTC) is formed in a double-membrane vesicle (Sawicki & Sawicki, 2005) by two large polyproteins (pp1a and pp1b), which encode non-structural proteins (Millet & Whittaker, 2015). The continuous replication by the RTC results in the formation of many sub-genomic RNAs (Hussain et al., 2005) which code for structural and auxiliary proteins. Virus assembly and budding takes place in smooth-walled vesicles in the endoplasmic reticulum, Golgi intermediate compartment (ERGIC) (Masters, 2006), and finally the virion-containing vesicles fuse with the plasma membrane to release the virus by exocytosis.

Many studies have established a strong link between the regulation of the innate immune response, the development of adaptive immunity, and the severity of COVID-19 (Mason, 2020; Mehrian-Shai, 2020; Tay et al., 2020). A hyper-inflammatory response was found in patients’ blood, nasopharyngeal samples, and bronchoalveolar lavage fluid, as evidenced by increased levels of cytokines such as IL-6, TNF-a, and MCP-1 that may lead to a severe acute respiratory syndrome (SARS), extensive coagulopathy, and multiorgan failure (Mason, 2020; Mehrian-Shai, 2020; Tay et al., 2020). Therefore, patients with severe COVID-19 require oxygen supplementation and intensive care, potentially exposing them to secondary opportunistic infections (Clancy & Nguyen, 2020; Friedland & Haribabu, 2020; Rawson et al., 2020; Ripa et al., 2021). As a result, current guidelines suggest the use of anticoagulant, anti-inflammatory, and antiviral medication along with broad-spectrum antibiotics and antifungals in patients with suspected or confirmed COVID-19 (Clancy & Nguyen, 2020; Friedland & Haribabu, 2020; Mehrian-Shai, 2020; Rawson et al., 2020). However, even with the same clinical intervention, patients display distinct trajectories with vastly different recovery times, clinical outcomes, or mortality (“Detrimental Effect of Diabetes and Hypertension on the Severity and Mortality of COVID-19 Infection: A Multi-Center Case-Control Study from India,” 2021). The molecular origin of such diverse outcomes are poorly understood in the context of lung pathophysiology, and only a handful of underpowered primary datasets have been published (Nienhold et al., 2020; Sanchez-Cerrillo et al., 2020; M. Wu et al., 2020; Xiong et al., 2020; Zhou et al., 2020).

This situation is further complicated by the emergence of coinfections in COVID-19 patients due to immunosuppression that may cause mucormycosis (Gupta et al., 2021; Moona & Islam, 2021; Prakash & Chakrabarti, 2021; Sen et al., 2021). Studies during previous SARS and MERS epidemics showed individuals receiving invasive mechanical ventilation were more likely to develop secondary infections and have a higher fatality rate (Feldman & Anderson, 2021). Recent investigations indicate that coinfections and/or superinfections occur at varying frequencies in oral, blood, and urine samples from COVID-19 patients (Alhumaid et al., 2021; Charalampous et al., 2020; Langford et al., 2020; Mostafa et al., 2020; Ripa et al., 2021; Rodriguez et al., 2021; Silva et al., 2021; Vijay et al., 2021). However, little is known about the prevalence of these pathogens and their exact molecular relevance in the human lung tissue during COVID-19 infection. Given that most COVID-19 deaths are due to pneumonia-related complications, it is critical to identify pathogens that co-infect severe COVID-19 patients and perform targeted therapeutic interventions.

In this work, we perform whole transcriptome sequencing of autopsy lung tissue from 31 patients who died due to severe COVID-19 related complications, and compared them to lung biopsies from 10 control patients who are not infected with SARS-CoV-2. Using metatranscriptomics, we determine characteristic changes to the host-transcriptome and unique microbial diversity in the lung parenchyma of severe COVID-19 patients. We map the host response at the level of genes, pathways, and change in cell-type abundances while identifying unique microbiome signatures driving dysbiosis in severe COVID-19 patients. Further, we correlate these findings with clinical features of the disease and dissect the potential molecular etiology of the disease that may help explain diverse outcomes leading to complications and suggest potential personalized therapeutic interventions.

## Methods

### Ethics Statement

The study followed the standards and principles established by India’s Directorate General of Health Services and Drug Controller General. Ethics approval was granted in writing from the Institute Ethics Committee of the All India Institute of Medical Sciences, New Delhi, India (IEC-538/05.06.2020, OP-28/05.02.2021). Consent was acquired from each patient’s personal and/or family members for autopsy, biopsy, and sample collection, in accordance with the Ethics approval.

### Patients and sample collection

This retrospective study examined 60 consecutive severe COVID-19 patients’ autopsies performed at the All India Institute of Medical Sciences in New Delhi, India, between September 2020 and December 2020 for patients who spent the last few days of their lives in the Intensive Care Unit (ICU). Minimally invasive post-mortem tissue sampling was performed in less than an hour on individuals with premortem PCR-confirmed SARS-CoV-2 infection at a biosafety level 3 post-mortem facility. The control (uninfected) lung samples (n = 10) represent healthy tissue taken from patients with lung cancer as part of standard medical evaluation during biopsy and/or surgical resection (>5cm from site of tumor from early stage Non-small cell lung cancer patients undergoing curative surgical resection). These control samples have been collected between August 2017 and July 2019. Paracancerous lung tissues have been taken as control for COVID-19 samples in other studies (Leng et al., 2020; S. Wang et al., 2021). Lung tissue was harvested from the parenchyma region in accordance with a standard protocol for histology and snap-frozen immediately, for RNA extraction. Lung tissues were fixed in 10% formalin, cut to the proper size and shape, embedded in paraffin for histological examination, or treated with TRIzol (Life Technologies), snap-frozen in liquid nitrogen, and kept at -80°C for RNA extraction. 31 out of 60 severe COVID-19 patient samples were chosen for analysis based on RNA yield and RNA quality (average RNA Integrity Number = 6.09). “Severe” COVID-19 was defined according to Ministry of Health and Family Welfare, Government of India guidelines (Government of India & Ministry of Health and Family Welfare, 2021). The guidelines define severe patients as the ones “characterized by a dysregulated immune response with hyperinflammation with subsequent development of ARDS”. These patients would be expected to have “acute respiratory infection with a history of fever or measured fever of ≥38 C°; and cough; with onset within the last 10 days; and requires hospitalization”. According to the definition of “severe” in the guidelines, the patient would present with severe pneumonia or acute respiratory distress syndrome or sepsis or septic shock. De-identified clinical information was extracted from patients medical records (Supplementary Table 1).

### Histopathological evaluation

Formalin-fixed, paraffin-embedded (FFPE) lung tissue blocks were processed and stained with hematoxylin and eosin using a standardized procedure. Two thoracic pathologists (DJ and AN) independently evaluated the slides. The following features: the extent of lung damage, injury, inflammation, presence or absence of hyaline membrane formation, lymphocyte infiltration, organizing pneumonia, alveolar fibrin deposition, fibrosis, and histologic features of type 2 pneumocyte hyperplasia, were noted and documented.

### RNA extraction, library preparation, and sequencing

Total RNA was extracted from lung tissue using the Maxwell automated instrument and the Maxwell® RSC Viral Total Nucleic Acid Purification Kit (Promega). The concentration of RNA and quality were measured with HS Total RNA 15nt.methods (Agilent) or Qubit RNA HS Assay (Thermofisher). The NGS library was prepared after cytoplasmic and mitochondrial rRNA depletion, using TruSeq Stranded Total RNA Gold kit per manufacturers’ instructions (Illumina, 20020598). The libraries were then sequenced on an Illumina NovaSeq6000 platform with 2×150 base pair reads (details of statistics given in Figure S1, Supplementary File 1).

### Host transcriptome analysis

Raw Illumina sequencing reads were checked for quality using FastQC (version 0.11.9)(*Babraham Bioinformatics - FastQC A Quality Control Tool for High Throughput Sequence Data*, n.d.) followed by adapter clipping and trimming using Trimmomatic (version 0.39)(Bolger et al., 2014) with default parameters. Trimmed reads were then aligned to the human reference genome (GRCh38, GENCODE v36)(Frankish et al., 2019; Schneider et al., 2017) using STAR aligner (version 2.7.8a)(Dobin et al., 2013). FeatureCounts (subread package version 2.0.1)(Y. Liao et al., 2014) was used to quantify gene expression. Quality checks were performed at each step using the MultiQC tool (version 1.10.1)(Ewels et al., 2016). Differential gene expression analysis was performed using the DESseq2 package (version 1.30.0)(Love et al., 2014) in R (version 4.0.3). The analysis was performed by removing the effects of confounding variables such as age and gender (Supplementary Table 1, Supplementary File 2) using the appropriate design formula. Genes with BH-adjusted p-value < 0.05 and absolute Log_2_ fold change greater than 1 in either direction were taken as significantly differentially expressed and Shrunken Log_2_ fold change values were used for further analysis. ClusterProfiler package (version 3.18.0)(Yu et al., 2012) was used for the Gene Ontology (GO) term Over Representation Analysis (ORA) of differentially expressed genes. GSVA package (version 1.38.2)(Hänzelmann et al., 2013) was used for all GSVA analysis and heatmaps of GSVA enrichment scores were visualized using the package pheatmap (version 1.0.12)(*Pheatmap: Pretty Heatmaps*, n.d.). Enriched gene sets had a corresponding change in GSVA enrichment scores with p < 0.05 using Wilcoxon test between the two groups compared. Boxplots and other visualizations were made using the ggplot2 package (version 3.3.3)(Wickham, 2011). All statistical tests were performed using functions from the base or stats package in R. For identification of transcription factors driving gene expression, we used the Enrichr tool (Kuleshov et al., 2016) using lists of genes upregulated in severe COVID-19 samples (and sub-groups as identified in the paper) when compared with controls.

### Curation of gene lists

Gene lists for GSVA were manually curated from various sources (Supplementary Table 2, Supplementary File 3). GSVA plots of Cell Types are based on published gene lists (Daamen et al., 2021). Gene lists for fibrosis and ECM (Extracellular Structure Organization) were taken from Wu *et al*., 2020 (M. Wu et al., 2020). Surfactant protein gene list was referenced from Islam and Khan, 2020(Islam & Khan, 2020). Gene lists from interferons, chemokines, interleukins, their receptors and other innate immune related pathways were sourced from HGNC (Vasiliou et al., 2021), ImmPort (Bhattacharya et al., 2018) and published gene lists (Daamen et al., 2021). KEGG pathways were utilized from MSigDB(Liberzon et al., 2011). Gene list for host proteins which interact with SARS-CoV-2 was referenced from Gordon *et al*., 2020 (Gordon et al., 2020).

### Cell deconvolution analysis

Multi-subject Single cell deconvolution (MuSic_0.2.0)(X. Wang et al., 2019) was used to predict the relative composition of different cell types from bulk RNA-Seq samples using existing single cell RNA-seq (scRNA-Seq) dataset as reference. From the relative composition of cell types, “Hedges’ g effect size” is measured using effsize_0.8.1 R package (Torchiano, 2016). Changes in cellular proportions comparing G1 vs N and G2 vs N were plotted for only those cell types that gave finite values in both G1 and G2, using a “scatterplot” using (ggplot2) (Wickham, 2011) R package.

### Connectivity Map (CMap) analysis

Connectivity Map (CMap)(Lamb et al., 2006) analysis was performed using the online portal https://clue.io/cmap to determine perturbagens (potential drugs reversing the aberrant gene expression) using the L1 version of CMap with L1000 data repository, Touchstone data set as a benchmark for assessing connectivity among perturbagens and Individual query option. The pertubagens were further filtered for potential therapeutic drugs.

### Metatranscriptomic analysis

Reads not mapped to the human genome were filtered to remove low complexity (entropy>=0.7), human rRNA and mitochondrial reads using BBMAP toolkit (version 38.90)(Bushnell, 2014). The filtered unmapped reads were then input into Seal (from the suite of bbtools) and binned into bacterial rRNA (using SILVA bacterial rRNA database)(Quast et al., 2013), human genome (GRCh38) and microbial bin. Taxonomic classification of reads was carried out on the microbial bin using Kraken2 (Wood et al., 2019) using organisms with at least 100 reads at the genus level for classification and confidence level of 0.3. The alpha diversity (Shannon diversity index) and bacterial taxon abundance was assessed using the PhyloSeq package (version 1.34.0)(McMurdie & Holmes, 2013).

### SARS-CoV-2 genomic and transcriptomic analysis

All COVID-19 samples with detectable SARS-CoV-2 reads were taken for further analysis. Filtered microbial reads from these samples were aligned against the SARS-CoV-2 reference genome(F. Wu et al., 2020) using BBMap (version 38.9)(Bushnell, 2014). Depth and coverage of the viral genome were obtained using samtools (version 1.9)(Li et al., 2009). Full length genomes were assembled for samples with high depth and coverage using SPAdes. The SARS-CoV-2 genomes were classified using GISAID (https://www.gisaid.org/), PANGO database (https://cov-lineages.org/) and nextclade (https://clades.nextstrain.org/) and were placed in a phylogenetic tree created using nextstrain (https://nextstrain.org/). Information on mutation types and frequency was obtained from http://giorgilab.unibo.it/coronannotator/. Additional information on strain B.1.36 was obtained from https://outbreak.info/situation-reports?pango=B.1.36. Transcriptome analysis was performed by aligning filtered viral reads to the reference strain (Wuhan-Hu-1) using Bowtie2 (Langmead & Salzberg, 2012). Read counts for the viral genes were obtained using featureCounts and normalized to Transcripts per Million (TPM) values.

### Data availability

All the data used in this study, which includes whole transcriptome sequencing raw data, filtered expression matrix that supports the findings of this study can be accessed in GEO under the accession number GSE183533. Further clinical annotation of the samples may be available upon reasonable request from the corresponding author DJ.

## Results

To characterize the pathology of SARS-CoV-2 infection, postmortem lung tissue samples were collected from 31 patients who had been diagnosed with severe COVID-19. As a control, 10 uninfected normal lung samples were taken from - patients diagnosed with cancer as part of standard surgery procedure (Figure 1). Among the 41 samples there were 19 (46%) females and 22 (54%) males. The 10 normal samples had 6 females and 4 males, while in the 31 COVID-19 samples there were 13 females and 18 males. The mean age for all 41 COVID-19 patients was 51.65 ± 15.27 years. The mean age for controls was 51 ± 16.22 years, while the COVID-19 patients were 57± 15.23 years (Table 1). Using nasopharyngeal swab PCR, all cases tested positive for SARS-CoV-2. At the time of onset, the most common symptoms were shortness of breath, fever, and cough.

**Figure 1.**
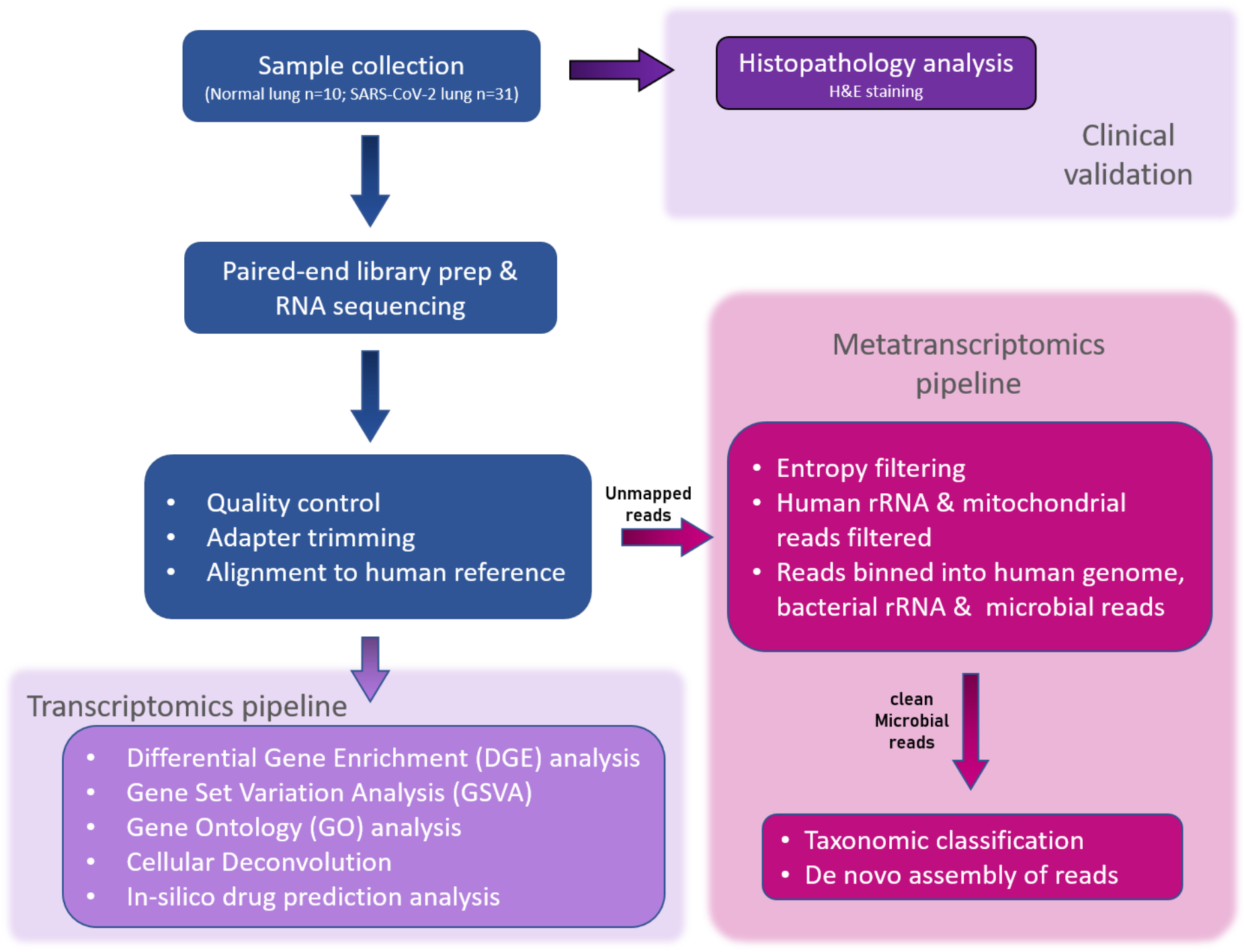
Schematic for histopathological and metatranscriptomic analysis pipeline. Lung tissue biopsy samples were collected from post mortem COVID-19 affected and control patients. The tissue samples were first evaluated for clinical physiology. The host rRNA was depleted prior to NGS library prep, the samples were then sequenced at an average depth of ∼47 million reads per sample. The sequence reads were filtered based on read quality. Filtered reads were used for differential gene expression, virome and microbiome profiling.

**Table 1.** Patient clinicopathological details.

| Characteristics | N or mean | % or range |
| --- | --- | --- |
| COVID-19 cohort (60 patients) |  |  |
| Age | 57y | 28-77y |
| Sex | M 18, F13 (1.4:1) |  |
| Hospitalisation time | 11.6d | 1-38d |
| Major comorbidities |  |  |
| Diabetes | 9 | 29.03% |
| Cardiovascular complications | 14 | 45.16% |
| Respiratory complications | 4 | 12.90% |
| Cancer | 5 | 16.12% |
| Liver or kidney complications | 15 | 48.38% |
| Thyroid complications | 3 | 9.67% |
| Patients with no prior comorbidities | 2 | 6.45% |
| Most common lung pathological features |  |  |
| Exudative phase of diffuse alveolar damage | 9 | 29.03% |
| Organizing phase diffuse alveolar damage | 6 | 19.35% |
| Acute bronchopneumonia | 3 | 9.67% |
| Co-existing acute bronchopneumonia with DAD | 5 | 16.12% |
| Moderate inflammatory changes | 5 | 16.12% |
| No changes/tissue sample | 3 | 9.67% |
| Control cohort (10 subjects) |  |  |
| Age | 51y | 21-69y |
| Sex | M 4, F6 (0.6:1) |  |
| Patients with smoking history | 6 |  |

The major cause of mortality was respiratory failure or multiorgan failure affecting the respiratory system. Consistent with a previous report (Carsana et al., 2020), fatal pulmonary tissue of all COVID-19 patients was highly abnormal, with evidence of diffuse alveolar damage (DAD), in addition to widespread hyaline membrane formation (the pathological hallmark of acute respiratory distress syndrome), acute lung injury, bronchopneumonia, and thrombosis being frequent (Supplementary Table 1, Supplementary File 2). COVID-19 afflicted patients’ lungs also show varying degrees of an inflammatory infiltrate.

To dissect the molecular aberrations underlying fatal COVID-19 cases, 41 lung tissue samples (31 cases and 10 control) were subject to whole transcriptome analysis using RNA-sequencing. The average number of reads ranged from 13.3 million to 115 million with an average of 46.9 million reads per sample. The range for the 10 normal samples is 35.4 million to 115 million with a mean of 58.9 million reads. While the range for 31 COVID-19 samples was slightly lower, at 13.3 million to 67.2 million with an average of 43.1 million reads, potentially owing to lung damage (Figure S1A, Supplementary File 1). The RNA-seq data was aligned to the human reference genome GRCh38 (gencode v36) to summarize gene counts. The alignment rate ranged from 60.0% to 89.6% with a mean of 83.72 ± 6.72% (Figure S1B, Supplementary File 1).

### Lung transcriptome signature in severe COVID-19

To interpret the host response, Differential Expression (DE) analysis was performed using the DESeq2 package in R accounting for confounding variables such as age and gender. Visualizing the gene level data per sample in a PCA plot we found that controls and a majority of COVID-19 samples clearly segregated on the first principal component (explaining 78% variance) except for 8 COVID-19 samples that clustered with normal samples (cov20, cov23, cov27, cov30, cov50, cov53, cov55, cov60). Henceforth, we refer to these 8 samples as G1 (group 1) and the rest of the COVID-19 samples as G2 (group 2), which can be seen colored differently in the PCA plot (Figure 2A). Next, we tried to check whether the gene expression pattern of G1 and G2 samples could be explained by the presence of any comorbidities. However, our analysis (Supplementary Table 1, Supplementary File 2) found no significant correlations (Figure S2 A-F, Supplementary File 1).

**Figure 2.**
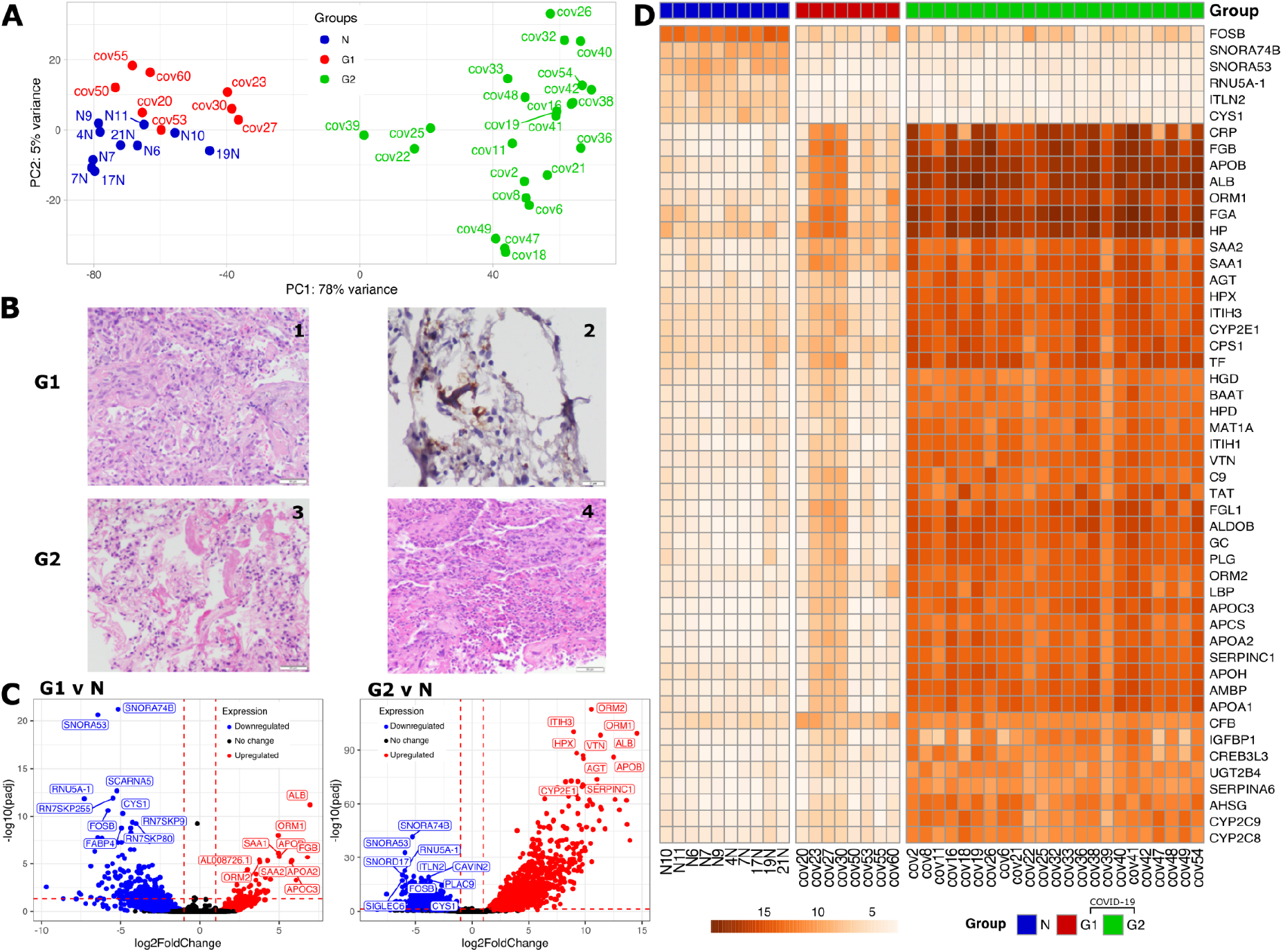
Differential expression and Histopathological analysis of COVID-19 samples. **A)** PCA plot based on gene expression counts across COVID-19 patients and Normal controls (blue) identified two groups of patients G1 (red) and G2 (green) samples **B)** Representative histological images of postmortem lung tissue sections for two patient groups. G1 group (top), displayed (**1**) acute organizing pneumonia (x200), (**2**) microthrombi (CD61 immunostain, x400). G2 group (below), displayed (**3**) diffuse alveolar damage with hyaline membranes (200x) and (**4**) acute bronchopneumonia with microabscesses (x200). **C)** Volcano plot describing the fold changes and FDR adjusted p-values between two groups of COVID-19 patients v/s Normal controls with down regulated genes in blue and upregulated genes in red. Top 10 DE gene names are highlighted. **D)**Variance-stabilizing transformed (vst) gene expression profile of top 50 significantly DE genes between control and two groups of COVID-19 patients.

We identified a total of 1,856 significantly differentially expressed genes (DEGs) between normal and all COVID-19 samples, with 864 genes significantly upregulated (Methods) and 992 genes significantly downregulated (Methods) in COVID-19 patients (Figure 2C). DE analysis between the G1 and Normal samples revealed the presence of 263 significant DEGs, with 56 genes significantly upregulated and 207 genes significantly downregulated, whereas, in the case of G2 group and normal samples, there were 3,094 DEGs, out of which 1,363 were significantly upregulated and 1,731 were significantly downregulated (Supplementary Table 3, Supplementary File 4). Between the two COVID-19 groups G2 and G1, there were 1,433 DEGs, out of which 1,314 were significantly upregulated and 119 were significantly downregulated (Figure 2C). Further differential expression analysis of G2 and G1 may be found in Supplementary Figure S7 (Supplementary File 1). A heatmap plotting the variance stabilized (vst) gene expression of the top 50 differentially expressed genes between all COVID-19 and normal samples clearly divides the samples into two groups and reaffirms the presence of two distinct categories of COVID-19 samples with gene expression of G1 matching that of normal samples (Figure 2D). 189 genes were differentially expressed in both G1 vs N and G2 vs N while 2899 genes were only differentially expressed in G2 vs N and 74 genes were only differentially expressed in G1 vs N (Supplementary Table 3, Supplementary File 4).

The most significantly upregulated genes in G1 and G2 include Orosomucoid 1 (ORM1 or better known as Alpha-1-acid glycoprotein 1 or AGP1), Orosomucoid 2 (ORM2 or Alpha-1-acid glycoprotein 2 or AGP2), Apolipoprotein B (APOB), ALB (Figure 2C, Figure 2D). Orosomucoid is known to be regulated by TNF beta, IL-1 beta, IL-6 and IL-6 related cytokines (Baumann et al., 1989; Fournier et al., 2000), along with immuno-modulating effects like inhibiting neutrophil migration (Mestriner et al., 2007) and has been employed as biomarker in COVID-19 (Shu et al., 2020). APOB has been observed to be upregulated in enterovirus 71 infection (Leong & Chow, 2006) and elevated levels in the blood of COVID-19 patients (Pushkarev et al., 2021). ALB levels in blood may be used as a biomarker for measuring COVID-19 severity (Liang et al., 2021).

Inter-Alpha-Trypsin Inhibitor Heavy Chain 3 (ITIH3), Hemopexin (HPX), Vitronectin (VTN), Angiotensinogen (AGT), SERPINC1 and CYP2E1 are certain genes that are highly upregulated in G2 when compared to normal samples (Figure 2C, Figure 2D). ITIH3 has been indicated as one of the plasma mortality markers for COVID-19 (Demichev et al., 2020; Völlmy et al., 2021). Angiotensinogen (AGT), is a hormone precursor involved in blood pressure regulation cascade that is implicated as potential biomarker and linked to severity of COVID-19 (Kouhpayeh et al., 2021; Sriram & Insel, 2020). VTN levels, from platelets, were observed to be high in case of SARS-CoV pneumonia as well (Lazzaroni et al., 2021). It has been noted that CYP2E1 has played a role in oxidative stress in Hepatitis C (Smirnova et al., 2016) and increased levels of the same have been associated with higher risk of adverse events, such as hepatotoxicity, especially in potential COVID-19 patients with obesity and Metabolic associated fatty liver disease (MAFLD) as a comorbidity (Ferron et al., 2020).

Some of the top 10 genes upregulated in G1 are Fibrinogen Beta Chain (FGB), Serum Amyloid A1 (SAA1), Serum Amyloid A2 (SAA2), Apolipoprotein A2 (APOA2), Apolipoprotein C3 (APOC3), AL008726.1 (Figure 2C). Increased abundance of Fibrinogen-β (FGB) also found in SARS-CoV-2 infected plasma exosomes is known to stimulate pro-inflammatory cytokine signaling (Sur et al., 2021). Proteomics studies showed the signatures of cytokine production and interferon-γ response, and increased level of SAA1 in the serum of COVID-19 patients (Singh et al., 2021). SAA2 may be a predictor of severity of COVID-19 (Papoutsoglou et al., 2021).

Small Nucleolar RNA, H/ACA Box 74B (SNORA74B), Small Nucleolar RNA, H/ACA Box 53 (SNORA53), RNU5A-1, FOSB and Cystin 1 (CYS1) are significantly downregulated in both G1 and G2, when compared to normal (Figure 2C, Figure 2D). Downregulation of CYS1 has been observed in another study for SARS-CoV, wherein low levels of CYS1 have been linked to activation of NF-kappa-B and subsequent cytokine storm (Zolfaghari Emameh et al., 2020). FOSB inactivation in mast cells has been shown to increase the inflammatory response (Duque-Wilckens et al., 2021) while contradictorily also reported to be upregulated in single cell analysis of CD4+ T cells of severe COVID-19 patients (Kalfaoglu et al., 2020).

SNORD17, ITLN2, CAVIN2, PLAC9 and SIGLEC6 are some of the genes highly downregulated in G2, along with SCARNA5, FABP4 and Pseudogenes (namely RN7SKP255, RN7SKP9, RN7SKP80) being highly downregulated in G1 (Figure 2C). Interlectin 2 (ITLN2) was found to be downregulated in a study of 22 blood samples of severe COVID-19 patients (Vastrad et al., 2020) as well, as seen in G2. Contrary to the trend observed in G2, in a single cell sequencing study of 16 COVID-19 patients, megakaryocyte progenitor cells/platelets showed increased expression of CAVIN2. SIGLEC6 belongs to family of Sialic acid binding immunoglobulin-like lectin proteins, out of which SIGLEC1, SIGLEC7 and SIGLEC10 have been implicated to play a role in COVID-19 (Doehn et al., 2021; Saheb Sharif-Askari et al., 2021). Low levels of Fatty acid-binding protein 4 (FABP4) in BALF macrophages of patients suffering from severe COVID-19, has been linked to declining lung function (M. Liao et al., 2020).

To identify biological processes implicated in the host response to SARS-CoV-2, we performed a Gene Ontology (GO) analysis of DEGs. The top 60 enriched GO terms (adjusted p-value < 0.05) were organized into a network of modules with edges connecting overlapping gene sets (Figure 3A). Key modules enriched in the G1 group included lipid metabolism, negative regulation of coagulation, and neutrophil-mediated immunity reported previously (Sanchez-Cerrillo et al., 2020; M. Wu et al., 2020; Xiong et al., 2020; Zhou et al., 2020). Key modules enriched in the G2 group were related to complement activation, xenobiotic metabolism, and peroxisomal protein transport as reported previously (Knoblach et al., 2021). No enriched modules were found downregulated in the G1 group while modules related to cilium formation, synapse formation, and membrane potential were downregulated in the G2 group suggesting suppression of neuronal processes as reported elsewhere (M. Wu et al., 2020).

**Figure 3.**
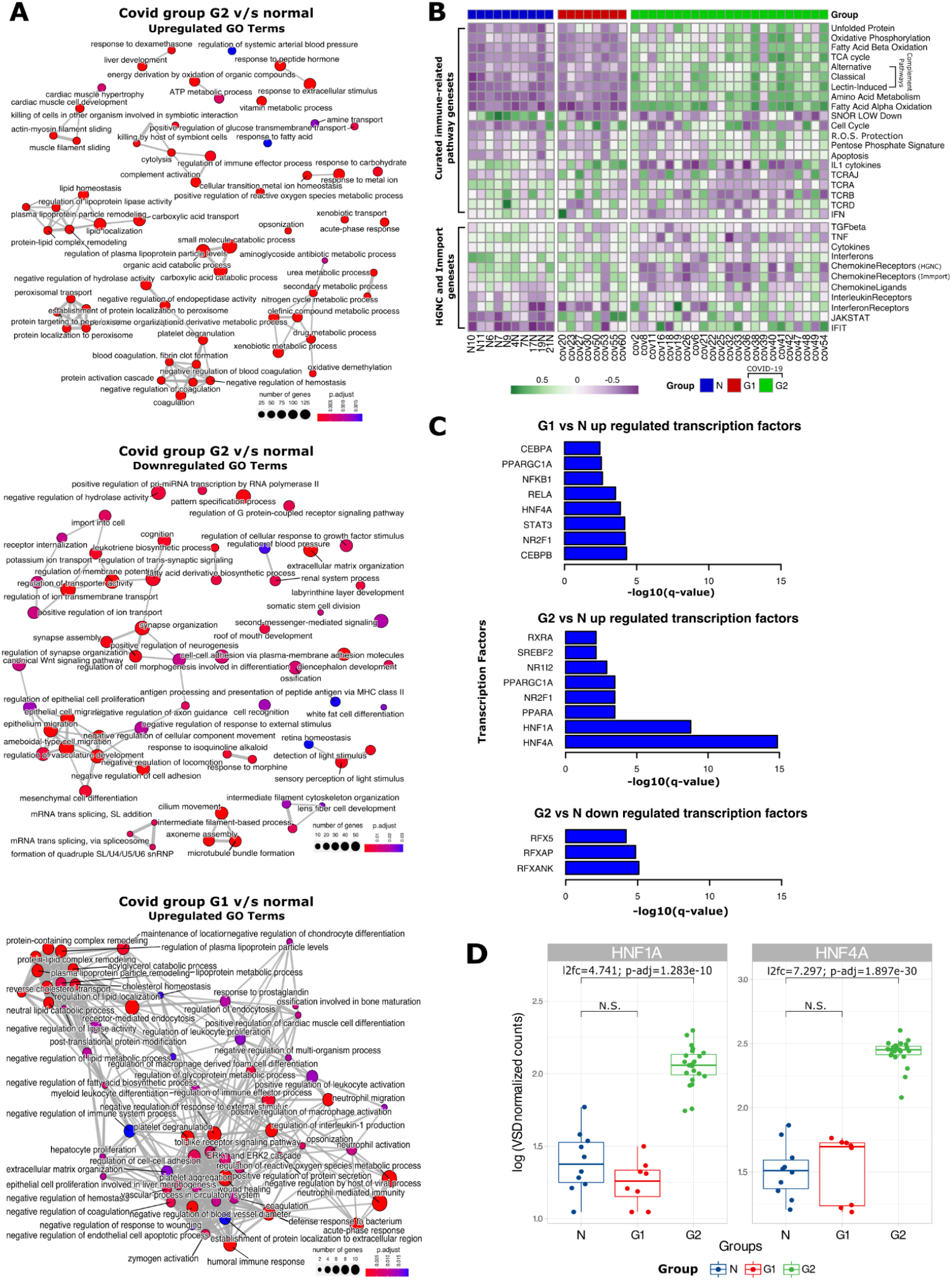
Gene Ontology enrichment, Gene Set Variation Analysis (GSVA) and Transcription Factor analysis between COVID-19 groups G1, G2 and normal control patients. **A)** GO enrichment map with nodes representing biological processes, edges representing overlapping gene sets. (top) up-regulated biological processes in G2 vs Normal individuals or ‘N’ (middle) down-regulated biological processes in G2 vs N (bottom) up-regulated biological processes in G1 vs N. **B)** Heatmap of GSVA depicts profiles of curated gene sets (Daamen et al., 2021),(Vasiliou et al., 2021),(Bhattacharya et al., 2018) across all the samples in our study. **C)** Barplots indicate the transcription factor targets enriched in significantly upregulated genes in G1 vs N, upregulated genes in G2 vs N, downregulated genes in G2 vs N. **D)** Boxplots of Log transformed vst-normalized gene counts representing differences among G1, G2 and normal sample groups for HNF1A and HNF4A

### Targeted gene set enrichment analysis using GSVA

To gain a better understanding of the regulation of key pathways identified in GO analysis we performed gene set variation analysis (GSVA). First, we analyze curated lists from HGNC and ImmPort (Supplementary Table 2, Supplementary File 3) of immune-related pathways, interferons, chemokines, interleukins and their receptors (*Gene Group*, n.d.). Expression of 11 of these gene lists was significantly altered with “Chemokine Ligands” altered only in G1 and 7 of the gene lists exclusively altered in G2 (Figure 3B). Genes corresponding to remaining 3 genelists namely, Interleukin receptors, Interferon Induced Transmembrane proteins (IFIT), and the JAK-STAT signaling were upregulated in both groups indicating a core antiviral inflammatory response (Sadler & Williams, 2008; Schoggins & Rice, 2011) in both patient groups. However, specific genes within each geneset were upregulated in each patient group suggesting distinct molecular pathways towards inflammation (Figure S3B, Supplementary File 1). Interleukin receptors IL20RA and IL5RA were upregulated in G1 while IL17RB, IL1RN, and IL22RA1 upregulated in the G2 group. Interleukins IL1RN and IL27 were only upregulated in G2, although IL6 was not upregulated in either groups, corroborating previous observations in lung tissue (M. Wu et al., 2020) despite high levels in blood of COVID-19 patients (Patel et al., 2021). Similarly, the entire chemokine ligand geneset itself was only upregulated in the G1 group, potentially due to high expression of chemokines such as CCL19 (Supplementary Figure S3E). Specific chemokines such as CCL16 were also upregulated in only the G2 group. We find further evidence of distinct inflammatory response within the G2 group through the exclusive downregulation of genes corresponding to Interferons, Chemokine receptors, Tumor Necrosis Factor mediated antiviral signaling, TGF beta signaling (Figure 3B); albeit with heterogeneity in expression of individual genes specially Bone Morphogenic Proteins (BMP), Growth Differentiation factors (GDF), and Inhibins (Figure S3B). Further, we find significant downregulation of genes associated with T-cell receptor signaling genes in the G2 group indicating T-cell dysfunction and potentially aberrant response.

Next, we dissected the molecular pathways that may be involved in the physical re-organization of lung tissue (Supplementary Table 2) using GSVA. We found that genes involved in Fibrosis, and Extracellular Structure Organization were significantly upregulated in both the patient groups (Figure 4A). Corroborating previous reports in COVID-19 patients (Islam & Khan, 2020), Surfactant proteins, which maintain surface tension in alveoli(Glasser & Mallampalli, 2012), were significantly downregulated. Again individual genes within each of these genesets displayed distinct expression between the two groups suggesting different routes towards aberrant lung physiology (Figure S3A, Supplementary File 1).

**Figure 4.**
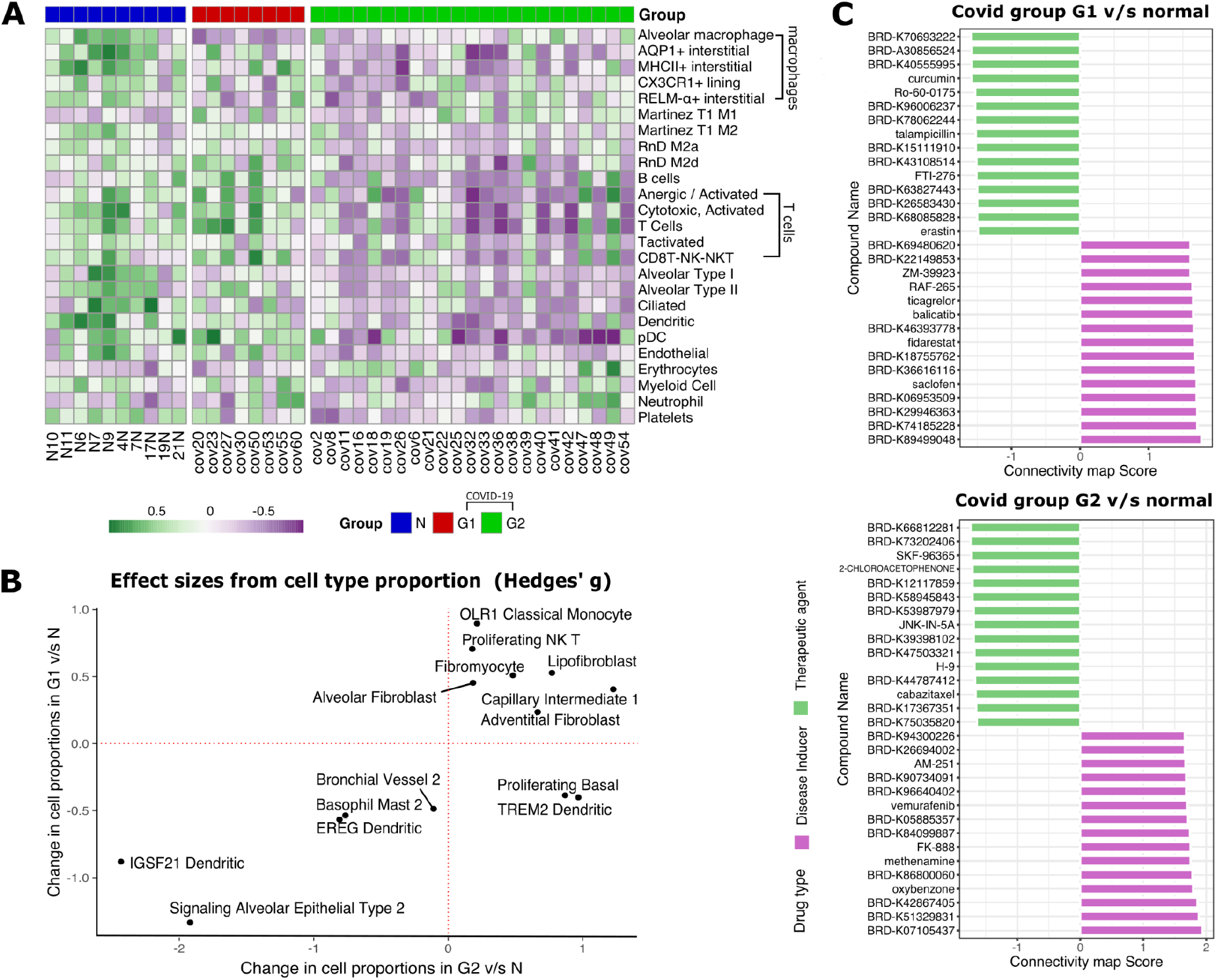
Cell-type specific gene-set enrichment and In-silico drug screening. **A)** Heatmap of per sample GSVA enrichment scores for previously implicated immune signaling pathways (above) and genes involved in physical organization of lung physiology (below) that are significantly differentially expressed (p<0.05, Wilcoxon test) between normal and either G1 or G2 samples. **B)** Changes in effective size of relative cellular proportions in G1 vs N and G2 vs N. Proliferating basal cells and TREM2+ dendritic cells were increased in G2 patients compared to G1. **C)** Bar plots of Connectivity Map (CMap) Scores for potential disease inducers (purple) and potential therapeutic compounds. (green) for G1 and G2 groups of patients respectively.

Next, we studied pathways previously found to be dysregulated in COVID-19. Out of a curated list of 36 pathways(Daamen et al., 2021), 20 (55.55%) pathways were found to be significantly upregulated between G1 or G2 relative to control samples. The G1 group showed significant overexpression of genes in cell cycle regulation and IL1 cytokines. While, the G2 group showed significant overexpression of genes related to apoptosis, all three complement pathways (classical, alternative, and lectin-induced) and metabolic pathways such as fatty acid oxidation, sugar metabolism, TCA cycle, and oxidative phosphorylation (Figure 3B). Massive change in metabolic pathways was further corroborated by GSVA using KEGG pathways whereby 85 out of 186 (45.69%) pathways were significantly altered (Figure S4A, Supplementary File 1) such as upregulation of steroid biosynthesis pathways and downregulation of neuroactive ligand-receptor interaction (corresponding to symptoms of anosmia and ageusia). Both the patient groups had characteristic dysregulation of TCR genes confirming previous bias(Gutierrez et al., 2020; P. Wang et al., 2021). For example in G1, TCRD was downregulated and TCRAJ was upregulated, while in G2, expression of both TCRA and TCRB was downregulated (Figure 3B).

We then studied host genes that directly interact with the viral proteins to help in viral entry and infection. The expression of both ACE2 and TMPRSS2, which play a role in SARS-CoV-2 entry(Hoffmann et al., 2020) did not differ between either group of patients when compared to normals (Figure S3D, Supplementary File 1). Cathepsins B and L which can substitute for TMPRSS2(Hoffmann et al., 2020) also remained unchanged (Figure S3D, Supplementary File 1). Next, we investigated the expression of 331 human proteins that directly bind to viral proteins(Gordon et al., 2020) in our samples (Supplementary Table 2, Supplementary File 3; Figure S3C, Supplementary File 1). Out of 248 (74.92%) highly expressed genes (depth greater than 100 reads in 90% of either normal or COVID-19 samples, hypothesizing that high expression would indicate a direct role), we found only 16 genes such as ERGIC1, GGH, PCSK6, PLOD2, that were differentially expressed only in the G2 group suggesting no significant over-representation the human host interactome in our patients (p = 0.985 using a hypergeometric test).

To find the regulators orchestrating the massive re-wiring of gene expression observed in COVID-19 patients, we investigated the enrichment of various targets of transcription factors (TFs) amongst our list of DEGs (Methods, Figure S5 A-C, Supplementary File 1). Targets of Hepatocyte nuclear factor-4 alpha (HNF-4α) and Nuclear respiratory factor 1 (NR2F1) were enriched in upregulated DEGs in both groups (Figure 3C). However, only HNF-4α was only overexpressed in the G2 patients when compared to Normal (Figure 3D) as well as G1 group (Supplementary Figure S7, Supplementary File 1). These results corroborate the role of HNF-4α in COVID-19 and other chronic lung pathologies (Agudelo et al., 2020; Nardacci et al., 2021), potentially through its role as a master regulator of lipid metabolism but only for the G2 group. This is further bolstered by lipid homeostasis related GO terms being enriched in upregulated genes of G2 when compared with G1. None of the other TF targets enriched in either upregulated genes or downregulated genes for both the groups were themselves expressed in the same direction as their targets - indicating alternate regulation.

### Cell-type deconvolution analysis maps altered lung cellular profile

Gene expression re-wiring can also be caused due to change in the proportion of different cell-types in the lung tissue of COVID-19 patients. To dissect this, first we performed GSVA using 57 gene expression signatures corresponding to different cell-types in the lung tissue(Daamen et al., 2021) and found that 25 signatures (43.86%) showed a significant change between COVID-19 patients and normal control patients, most of them being downregulated in G2 patients (Figure 4A). The only common signature between G1 and G2 groups was the loss of alveolar macrophages as reported previously (M. Liao et al., 2020). The G2 group showed an enrichment of signatures corresponding to Erythrocytes, damage-associated M1 macrophages and Neutrophils suggesting acute inflammation and structural damage. The G2 group also showed loss of signature corresponding to cells in the lung parenchyma such as alveolar epithelial cells, endothelial cells, and ciliated cells alongside loss of platelets suggesting prolonged lung damage and thrombocytopenia. Such reorganization of lung tissue was also accompanied by loss in signature corresponding to specific Lymphocytes such as B-cells, activated T-cells, NK-T-cells and anergic T-cells alongside loss of dendritic and plasmacytoid DCs (pDC) suggesting immune exhaustion.

To further dissect the finer details of lung physiology, we performed cell-type deconvolution to estimate the proportion of 58 different cell types using a published single-cell RNA-sequencing dataset from lungs (Travaglini et al., 2020) (Figure S4B, Supplementary File 1). In both the patient groups, we found a strong depletion in the proportion of IGSF21+ & EREG DCs, signaling Alveolar epithelial cells, and “Basophil Mast 2” cells alongside enrichment in the proportion of Lipofirboblasts, indicative of common pathophysiology. In the G1 group, we found a high proportion of proliferating NK-T cells and “OLR1 Classical Monocytes” alongside depletion of “Bronchial Vessel 2” cells. Interestingly, two cell populations showed antagonistic change in cell proportions namely Proliferating Basal and TREM2^+^ DCs that were depleted in G1 but enriched in the G2 group of patients (Figure 4B).

### In-silico drug screening using Connectivity maps

We performed connectivity map (CMap) analysis with differentially expressed genes from G1 vs Normal and G2 vs Normal samples to help predict the different set of drugs which can be used to reverse the molecular signature in the two types of patients. We identified 423,422 perturbagens and among them 136,460 were drug perturbagens which contain both therapeutic agents and inducers. We filtered them for compound data and A549 cell lines (adenocarcinomic human alveolar basal epithelial cells) and found 11456 unique drug perturbagens for both groups. Among these, 1,986 and 2,048 perturbagens had negative connectivity scores for G1 and G2 groups respectively, implying their potential as therapeutic agents that can be screened further using in-vivo experiments (Supplementary Table 4, Supplementary File 5) to identify potential drugs. Among the top therapeutic agents we found curcumin, Ro-60-0175, talampicillin, FTI-276, erastin as potential therapeutic agents for G1 patients while we found SKF-96365, 2-chloroacetophenone, JNK-IN-5A, H-9, cabazitaxel as potential therapeutic agents for G2 patients (Figure 4C).

### Metatranscriptome analysis reveals difference in species richness and distribution between control and COVID-19 lungs

To identify the microbial signature in patient lung tissue we performed a metatranscriptomic analysis (Figure 1). Briefly, reads that did not align to the human genome (2.23 ± 0.46% ; Figure S1C, Supplementary File 1) were filtered for low complexity sequences (Figure S1D, Supplementary File 1) and bacterial rRNA, and then used for k-mer based phylogenetic classification (Supplementary Table 5, Supplementary File 6). After taxonomic assignment, we found significant loss in species richness in both G1 (Wilcoxon test; p-value = 4.6e-05) and G2 (p=8.6e-08) patients compared to normal controls, with no difference between the two groups (Figure 5A). At the phylum level (Figure 5B), Actinobacteria were found to be less abundant in both G1 and G2 samples compared to normal controls (p=6.2e-04 and p=1.5e-0 respectively). While only in G2 patients, we found depletion of Proteobacteria (p=4.2e-06) and enrichment of Firmicutes (p-value=7.9e-06) (Figure S6A, Supplementary File 1).

**Figure 5.**
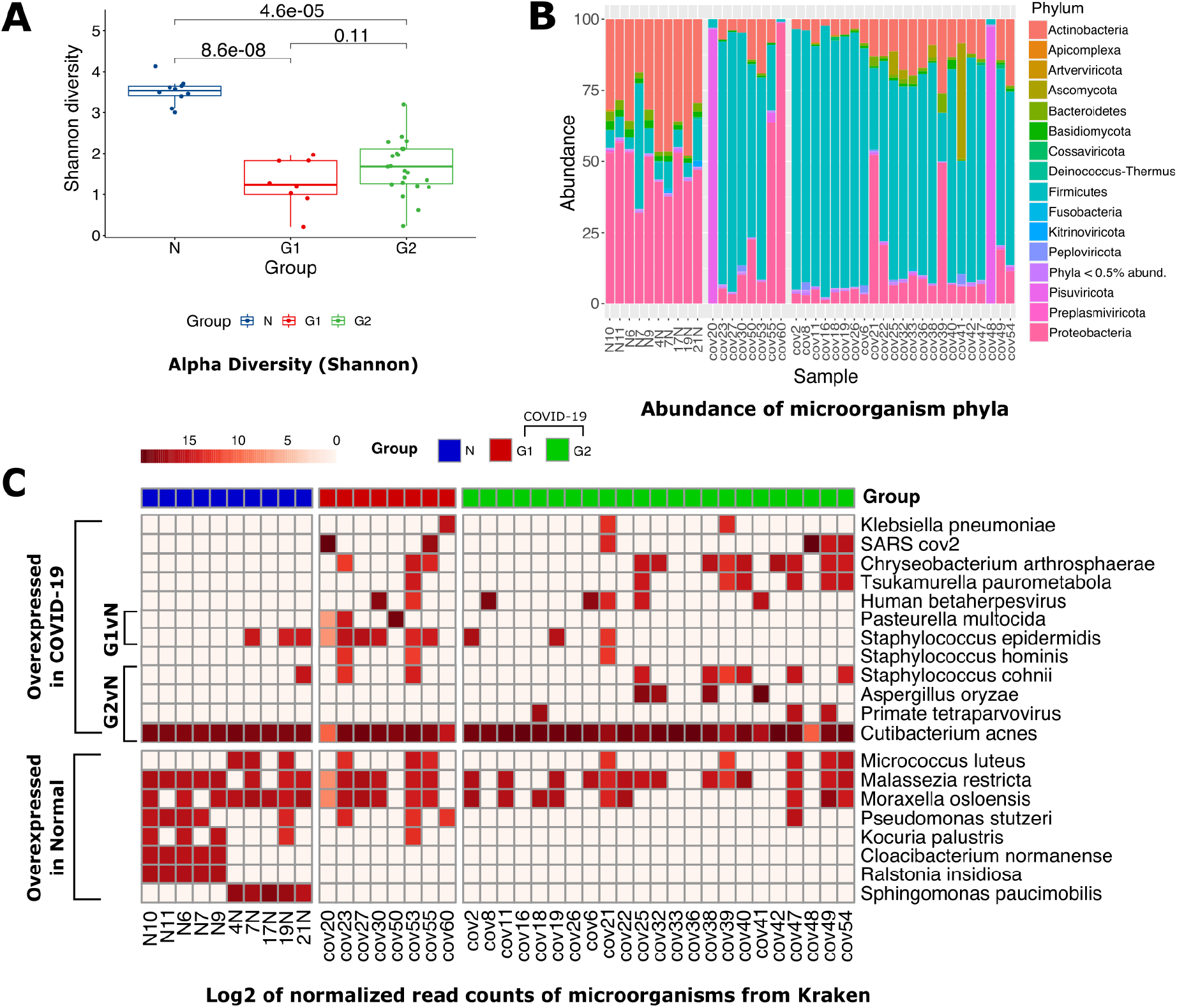
Lung microbiome of COVID-19 and control patients. **A)** Box Plots showing alpha diversity (Shannon index) for N, G1, and G2 groups **B)** Stacked barplot representing the distribution of bacterial taxon at the phylum level. **C)** Log2 of normalized read counts for the bacterial species present in at least 3 samples and which were present in both G1 and G2 as compared to the control group.

To look at changes in composition between COVID-19 patient groups at the species level and reduce noise, we opted to include only those species with a minimum of 100 reads across at least three samples. We found 20 species distributed between the three groups of patients based on normalized counts per million (CPM) values. We found that some *Staphylococcus* species such as *S. cohnii* were enriched in both groups of COVID-19 patients while other *Staphylococcus* species, such as *S. hominis, S. epidermidis* and *S. warneri*, along with multidrug resistant *Chryseobacterium arthrosphaerae* were enriched in the G1 group. Additionally, the G1 patients were also enriched for the pathogenic *Pasteurella multocida* while the G2 patients were enriched in other pathogens such as *Klebsiella pneumoniae* and *Tsukamurella paurometabola* (Figure 5C).

### Analysis of the co-isolated SARS-CoV-2 genome and transcriptome

After phylogenetic classification of filtered microbial reads, only 6 out of the 31 (19.35%) COVID-19 samples were found to have more than 100 reads assigned to SARS-CoV-2 (Supplementary table 6). Three samples with greater than 10X depth and above 99% coverage were used for de novo genome assembly using SPAdes to obtain a single contig of length > 29 kbp (corresponding to the full length SARS-CoV-2 genome) from two samples and 11 contigs spanning the entire genome were assembled from the third (deposited at GISAID accession ID : EPI_ISL_4392854,EPI_ISL_4392853,EPI_ISL_4392851).

Consensus calling was used for variant calling by aligning to the Wuhan-Hu-1 reference strain (Methods). Phylogenetic analyses assigned these sequences to the GISAID clade GH or Nextstrain clade 20A in PANGO lineage B.1.36 (Figure 6A) circulating in Europe, Asia and North America from September 2020 to March 2021 (Figure S6C, Supplementary File 1). Genomic sequence analysis revealed that cov20 had 12, cov48 had 13 while cov55 had 14 mutations with respect to the reference with 7 mutations were in the spike protein region. Most of the mutations were SNP’s (56.41 %) while C > T was the most frequently observed substitution (58.97%) (Figure 6B; Supplementary Table 6).

**Figure 6.**
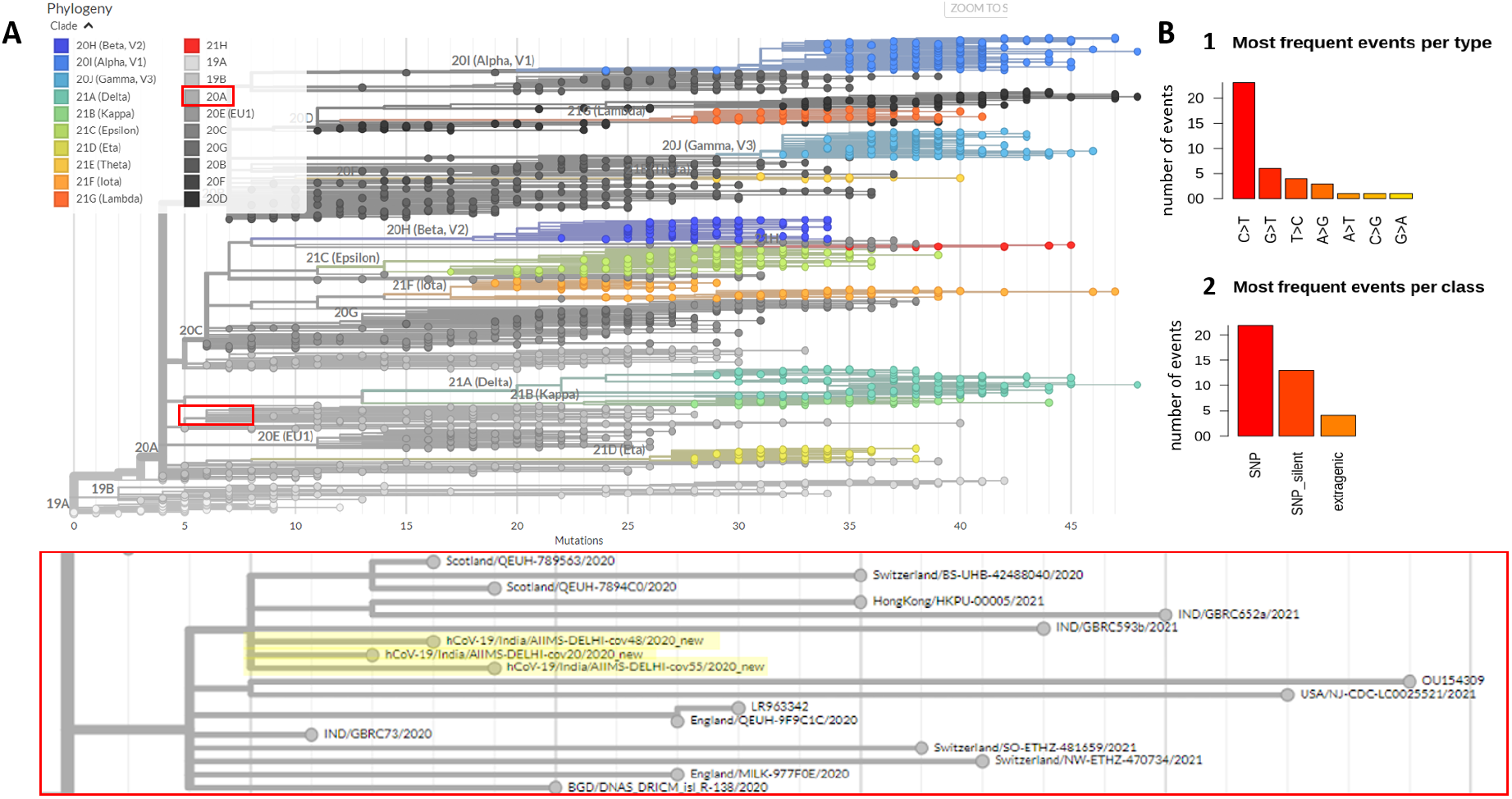
Phylogenetic classification and analysis of recovered SARS-CoV-2 genomes. **A)**. Phylogenetic analysis of the three high confidence SARS-CoV-2 genomes show that they all fall within the same nextstrain clade 20A and are next to each other in the phylogenetic tree. **B)**. Analysis of the mutations from all three genomes show that non synonymous SNPs are the most common mutation type observed while C > T base change was the most common mutation.

Given the depth of sequencing of SARS-CoV-2 virus in the three samples above, we were also able to estimate viral gene expression (Figure S6B, Supplementary File 1). We found that ORF10 had the highest expression level (mean TPM of 3.44 million) which was twice more than any other gene and is consistent with another report (Zhang et al., 2020). ORF7b had the lowest expression (mean TPM of 0.12 million), while the viral N gene, encoding the Nucleocapsid and a common target for RT-PCR diagnostic tests, had the second highest expression (mean TPM of 1.64 million). Since normal samples are collected much before COVID-19 pandemic (Methods) and sequenced in the same batch of COVID-19, very few reads of COVID-19 appear in our normal samples, which are most probably a contamination during library preparation and have been filtered out by setting higher detection threshold for detecting COVID-19 in samples to at least 100 reads.

## Discussion

Dissecting the molecular biology of primary lung tissue is essential for understanding drivers of mortality in severe COVID-19 infection to develop precision novel therapies and monitor disease prognosis. However, most large-scale cohort studies have been limited to non-invasive sampling from blood, nasopharyngeal fluid, and bronchoalveolar lavage to delineate the effects of the disease in peripheral tissues (Sanchez-Cerrillo et al., 2020; M. Wu et al., 2020; Xiong et al., 2020; Zhou et al., 2020) that consider all severe patients as a singular group. Most studies on lung tissue from severe cases of COVID-19 profiled either FFPE samples with compromised RNA or were underpowered with less than 18 patients (Nienhold et al., 2020; Sanchez-Cerrillo et al., 2020; M. Wu et al., 2020; Xiong et al., 2020; Zhou et al., 2020). Here, we describe the molecular pathology of severe COVID-19 patients in the largest cohort lung tissue biopsies from 31 post-mortem patients compared with biopsies of non-cancerous tissue from 10 lung cancer patients as a control group. We found that COVID-19 patient lung tissue displayed two distinct molecular signatures defined by the lung transcriptome profiles. The lung transcriptome did not show indication of bacterial or fungal co-infections in severe COVID-19 patients.

The dominant “classical” signature found in 74% of patients displayed a large-scale reorganization of gene expression characterized by loss in various parenchymal cells, unfolded protein response, enhanced complement system supported by metabolic reprogramming, Neutrophil upregulation, and activated T-cells depletion. The rarer “Cytokine Release Syndrome” (CRS) signature found in 26% of patients showed minimal deviation in gene expression from normal controls but was marked by the proliferation of NK-T-cells, and enhanced production of cytokines such as IL1 and CCL19. A limitation of the comparisons drawn for significantly downregulated genes for COVID-19 would be that those genes are simply upregulated in cancer-adjacent normal tissues (due to them being upregulated in cancer) and is acknowledged as thus. TREM2+ DCs, which play a role in T-cell priming and are found in the BALF of severe COVID-19 patients(M. Liao et al., 2020), were enriched in the classical signature but depleted in the CRS signature. Damage-associated M1 macrophage signature was enhanced in the dominant group and was depleted in the rare group while the reverse pattern was seen for the protective M2 macrophages. Although a previous study by Nienhold et al (Nienhold et al., 2020) also described two distinct subtypes of COVID-19 based on the expression of Interferon stimulated genes (ISG); the two patient subtypes identified in this study could not be segregated based on ISG expression.

We propose two models to explain the disease signatures based on complement activation and failure to launch an adaptive immune response. The dominant signature represents “classic” COVID-19 patients with an acute infection due to high initial viral load, as evidenced by sustained unfolded protein response (Chan, 2014), leading to hyperinflammation through the complement system activating neutrophils. This hyperinflammation leads to lung damage (Figure 2B) and recruitment of M1 macrophages. Also, these patients are unable to mount an adaptive immune response as evidenced by the depletion of T-cells, potentially due to direct viral infection, or T-cell exhaustion due to alternate metabolic flux. The rarer signature may represent patients with cytokine release syndrome (Henderson et al., 2020; Hu et al., 2021; Tang et al., 2020) at low viral load, who are unable to deploy the complement system and depend on innate immunity through NK-T-cells. These patients may have characteristic “lung damage”, repaired by M2 macrophages. However, these patients too are unable to mount an adaptive immune response due to lymphopenia or diminished T-cell priming as a result of depleted TREM2^+^ DCs.

Although both groups of COVID-19 patients were given the same treatment, they displayed differences in the infection spectrum. Patients with the rare signature were specifically enriched for *Staphylococcus epidermidis* and *Pasteurella multocida* despite administration of broad spectrum antibiotics such as Azithromycin. Effect of broad spectrum antibiotics can be perceived from the reduced species diversity in severe COVID-19 patients. Further, we found that the rare signature patients may have impaired immunity to deal with such infections through downregulation of genes involved in CCR6 dependent bactericidal activity (DEFB1) and the downregulation of de novo biosynthesis of steroids from cholesterol (HSD17B6, GREM2, FADS6, AADAC). These characteristic changes in the lung microbiome warrant further investigation into the contribution of nasopharyngeal and airway microbiota in COVID-19 patients with lung complications.

It must be noted that the microbiome of lung tissue collected during aseptic surgery of cancer patients will significantly differ from postmortem microbiome of COVID-19 patients. Regardless, our results indicate that there is a dysbiosis in lung tissue of patients with severe COVID-19 reinforced by the fact that lung tissues extracted in sterile conditions (Normal samples) have more alpha diversity than the COVID-19 postmortem lung tissues.

Despite most patients clearing the SARS-CoV-2 virus, we were able to recover the full genome from 3 patients that converged to the same consensus indicating clonal viral expansion. Upon analyzing the SARS-CoV-2 genome, we found that the ORF10 non-coding gene, with a low mutation rate and no selection pressure (Nguyen et al., 2021), had the highest gene expression with twice the number of transcripts as compared to the N gene, suggesting its potential use as a novel target for RT-PCR testing. Also the 3 genomes recovered are from the same lineage (Figure 6). With 3 genomes, one cannot significantly conclude whether these strains have led to different trajectories in the two groups of patients and if we had been able to retrieve complete genome sequences for more samples, we could interpret the cause in accordance with the strain of the virus.

In conclusion, our work supports further clinical investigation correlating prognosis by stratifying patients based on the circulating molecules involved in complement activation shown recently (Ma et al., 2021), or characteristic cytokines such as CCL19, implicated in COVID-19 mortality (Balnis et al., 2021). Such non-invasive stratification of patients can be used to test the efficacy drugs identified in our study that reverse the molecular changes for the two patient signatures such as cabazitaxel to treat “classical” patients or talampicillin to treat “CRS” patients.

This is the first study analyzing the lung transcriptome in the Indian population, one of the worst affected countries in the world with over 400,000 COVID-19 mortalities. Although about 1 in 7 people in the world comes from India, data from Indian populations is often missing from such molecular investigations of diseases and COVID-19 is no different. Our study bridges the gap of diversity in sampling various populations across the world that are affected by the pandemic. This is particularly of interest as in our recent work we demonstrate a significant contribution of genetics towards mortality in severe COVID-19 (Prakrithi et al., 2021). Therefore, we conclude by hypothesizing that the heterogeneity in the molecular signature of severe COVID-19 patients may be driven by patient genetics and can be used for candidate gene prioritization.

## Supplementary Files

1. Supplementary File 1 (Supplementary figures)
2. Supplementary File 2 (Table S1)
3. Supplementary File 3 (Table S2)
4. Supplementary File 4 (Table S3)
5. Supplementary File 5 (Table S4)
6. Supplementary File 6 (Table S5)
7. Supplementary File 7 (Table S6)

## References

Agudelo, C. W., Samaha, G., & Garcia-Arcos, I. (2020). Alveolar lipids in pulmonary disease. A review. Lipids in Health and Disease, 19(1), 122.

Ahmadpoor, P., & Rostaing, L. (2020). Why the immune system fails to mount an adaptive immune response to a COVID-19 infection [Review of Why the immune system fails to mount an adaptive immune response to a COVID-19 infection]. Transplant International: Official Journal of the European Society for Organ Transplantation, 33(7), 824–825.

Alhumaid, S., Al Mutair, A., Al Alawi, Z., Alshawi, A. M., Alomran, S. A., Almuhanna, M. S., Almuslim, A. A., Bu Shafia, A. H., Alotaibi, A. M., Ahmed, G. Y., Rabaan, A. A., Al-Tawfiq, J. A., & Al-Omari, A. (2021). Coinfections with Bacteria, Fungi, and Respiratory Viruses in Patients with SARS-CoV-2: A Systematic Review and Meta-Analysis. Pathogens, 10(7). https://doi.org/10.3390/pathogens10070809

Babraham bioinformatics - FastQC A quality control tool for high throughput sequence data. (n.d.). Retrieved August 20, 2021, from https://www.bioinformatics.babraham.ac.uk/projects/fastqc/

Balnis, J., Adam, A. P., Chopra, A., Chieng, H. C., Drake, L. A., Martino, N., Bossardi Ramos, R., Feustel, P. J., Overmyer, K. A., Shishkova, E., Coon, J. J., Singer, H. A., Judson, M. A., & Jaitovich, A. (2021). Unique inflammatory profile is associated with higher SARS-CoV-2 acute respiratory distress syndrome (ARDS) mortality. American Journal of Physiology. Regulatory, Integrative and Comparative Physiology, 320(3), R250–R257.

Baumann, H., Prowse, K. R., Marinković, S., Won, K. A., & Jahreis, G. P. (1989). Stimulation of hepatic acute phase response by cytokines and glucocorticoids. Annals of the New York Academy of Sciences, 557, 280–295, discussion 295–296.

Bhattacharya, S., Dunn, P., Thomas, C. G., Smith, B., Schaefer, H., Chen, J., Hu, Z., Zalocusky, K. A., Shankar, R. D., Shen-Orr, S. S., Thomson, E., Wiser, J., & Butte, A. J. (2018). ImmPort, toward repurposing of open access immunological assay data for translational and clinical research. Scientific Data, 5, 180015.

Bolger, A. M., Lohse, M., & Usadel, B. (2014). Trimmomatic: a flexible trimmer for Illumina sequence data. Bioinformatics, 30(15), 2114–2120.

Bushnell, B. (2014). BBMap: A Fast, Accurate, Splice-Aware Aligner. https://escholarship.org/uc/item/1h3515gn

Carsana, L., Sonzogni, A., Nasr, A., Rossi, R. S., Pellegrinelli, A., Zerbi, P., Rech, R., Colombo, R., Antinori, S., Corbellino, M., Galli, M., Catena, E., Tosoni, A., Gianatti, A., & Nebuloni, M. (2020). Pulmonary post-mortem findings in a series of COVID-19 cases from northern Italy: a two-centre descriptive study. The Lancet Infectious Diseases, 20(10), 1135–1140.

Chan, S.-W. (2014). The unfolded protein response in virus infections. Frontiers in Microbiology, 5, 518.

Charalampous, T., Alcolea-Medina, A., Snell, L. B., Williams, T. G. S., Batra, R., Camporota, L., Meadows, C. I. S., Wyncoll, D., Barrett, N. A., Hemsley, C. J., Bryan, L., Newsholme, W., Boyd, S. E., Green, A., Mahadeva, U., Patel, A., Cliff, P. R., Page, A. J., O’Grady, J., & Edgeworth, J. D. (2020). Application of respiratory metagenomics for COVID-19 patients on the intensive care unit to inform appropriate initial antimicrobial treatment and rapid detection of nosocomial transmission. medRxiv, 2020.11.26.20229989.

Clancy, C. J., & Nguyen, M. H. (2020). Coronavirus Disease 2019, Superinfections, and Antimicrobial Development: What Can We Expect? Clinical Infectious Diseases: An Official Publication of the Infectious Diseases Society of America, 71(10), 2736–2743.

Daamen, A. R., Bachali, P., Owen, K. A., Kingsmore, K. M., Hubbard, E. L., Labonte, A. C., Robl, R., Shrotri, S., Grammer, A. C., & Lipsky, P. E. (2021). Comprehensive transcriptomic analysis of COVID-19 blood, lung, and airway. Scientific Reports, 11(1), 7052.

Demichev, V., Tober-Lau, P., Nazarenko, T., Thibeault, C., Whitwell, H., Lemke, O., Röhl, A., Freiwald, A., Szyrwiel, L., Ludwig, D., Correia-Melo, C., Helbig, E. T., Stubbemann, P., Grüning, N.-M., Blyuss, O., Vernardis, S., White, M., Messner, C. B., Joannidis, M., … PA-COVID-19 Study group. (2020). A time-resolved proteomic and diagnostic map characterizes COVID-19 disease progression and predicts outcome. In bioRxiv. medRxiv. https://doi.org/10.1101/2020.11.09.20228015

Detrimental effect of diabetes and hypertension on the severity and mortality of COVID-19 infection: A multi-center case-control study from India. (2021). Diabetes & Metabolic Syndrome: Clinical Research & Reviews, 15(5), 102248.

Dobin, A., Davis, C. A., Schlesinger, F., Drenkow, J., Zaleski, C., Jha, S., Batut, P., Chaisson, M., & Gingeras, T. R. (2013). STAR: ultrafast universal RNA-seq aligner. Bioinformatics, 29(1), 15–21.

Doehn, J.-M., Tabeling, C., Biesen, R., Saccomanno, J., Madlung, E., Pappe, E., Gabriel, F., Kurth, F., Meisel, C., Corman, V. M., Hanitsch, L. G., Treskatsch, S., Heim, K., Stegemann, M. S., Ruwwe-Glösenkamp, C., Müller-Redetzky, H. C., Uhrig, A., Somasundaram, R., Spies, C., … Hübner, R.-H. (2021). CD169/SIGLEC1 is expressed on circulating monocytes in COVID-19 and expression levels are associated with disease severity. Infection, 49(4), 757–762.

Dong, E., Du, H., & Gardner, L. (2020). An interactive web-based dashboard to track COVID-19 in real time. The Lancet Infectious Diseases, 20(5), 533–534.

Duque-Wilckens, N., Maradiaga, N. C., Szu-ying, Y., Nestler, E. J., Robison, A. J., & Moeser, A. J. (2021). Mast cell-specific inactivation of Fosb exacerbates release of pro-inflammatory mediators in models of systemic anaphylaxis and lipopolysaccharide-induced sepsis. The Journal of Immunology, 206(1 Supplement), 97.12–97.12.

Ewels, P., Magnusson, M., Lundin, S., & Käller, M. (2016). MultiQC: summarize analysis results for multiple tools and samples in a single report. Bioinformatics, 32(19), 3047–3048.

Feldman, C., & Anderson, R. (2021). The role of co-infections and secondary infections in patients with COVID-19. Pneumonia (Nathan Qld.), 13(1), 5.

Ferron, P.-J., Gicquel, T., Mégarbane, B., Clément, B., & Fromenty, B. (2020). Treatments in Covid-19 patients with pre-existing metabolic dysfunction-associated fatty liver disease: A potential threat for drug-induced liver injury? Biochimie, 179, 266–274.

Fournier, T., Medjoubi-N, N., & Porquet, D. (2000). Alpha-1-acid glycoprotein. Biochimica et Biophysica Acta, 1482(1-2), 157–171.

Frankish, A., Diekhans, M., Ferreira, A.-M., Johnson, R., Jungreis, I., Loveland, J., Mudge, J. M., Sisu, C., Wright, J., Armstrong, J., Barnes, I., Berry, A., Bignell, A., Carbonell Sala, S., Chrast, J., Cunningham, F., Di Domenico, T., Donaldson, S., Fiddes, I. T., … Flicek, P. (2019). GENCODE reference annotation for the human and mouse genomes. Nucleic Acids Research, 47(D1), D766–D773.

Friedland, R. P., & Haribabu, B. (2020). The role for the metagenome in the pathogenesis of COVID-19. EBioMedicine, 61, 103019.

Gene group. (n.d.). Retrieved July 31, 2021, from https://www.genenames.org/data/genegroup/#!/

Glasser, J. R., & Mallampalli, R. K. (2012). Surfactant and its role in the pathobiology of pulmonary infection. Microbes and Infection / Institut Pasteur, 14(1), 17–25.

Gordon, D. E., Jang, G. M., Bouhaddou, M., Xu, J., Obernier, K., White, K. M., O’Meara, M. J., Rezelj, V. V., Guo, J. Z., Swaney, D. L., Tummino, T. A., Hüttenhain, R., Kaake, R. M., Richards, A. L., Tutuncuoglu, B., Foussard, H., Batra, J., Haas, K., Modak, M., … Krogan, N. J. (2020). A SARS-CoV-2 protein interaction map reveals targets for drug repurposing. Nature, 583(7816), 459–468.

Government of India, & Ministry of Health and Family Welfare. (2021). Updated Detailed Clinical Management Protocol for COVID19 adults dated 24 05 2021. https://www.mohfw.gov.in/pdf/UpdatedDetailedClinicalManagementProtocolforCOVID19adultsdated24052021.pdf

Gupta, A., Sharma, A., & Chakrabarti, A. (2021). The emergence of post-COVID-19 mucormycosis in India: Can we prevent it? Indian Journal of Ophthalmology, 69(7), 1645–1647.

Gutierrez, L., Beckford, J., & Alachkar, H. (2020). Deciphering the TCR Repertoire to Solve the COVID-19 Mystery. Trends in Pharmacological Sciences, 41(8), 518–530.

Hänzelmann, S., Castelo, R., & Guinney, J. (2013). GSVA: gene set variation analysis for microarray and RNA-seq data. BMC Bioinformatics, 14, 7.

Henderson, L. A., Canna, S. W., Schulert, G. S., Volpi, S., Lee, P. Y., Kernan, K. F., Caricchio, R., Mahmud, S., Hazen, M. M., Halyabar, O., Hoyt, K. J., Han, J., Grom, A. A., Gattorno, M., Ravelli, A., De Benedetti, F., Behrens, E. M., Cron, R. Q., & Nigrovic, P. A. (2020). On the Alert for Cytokine Storm: Immunopathology in COVID-19. Arthritis & Rheumatology (Hoboken, N.J.), 72(7), 1059–1063.

Hoffmann, M., Kleine-Weber, H., Schroeder, S., Krüger, N., Herrler, T., Erichsen, S., Schiergens, T. S., Herrler, G., Wu, N.-H., Nitsche, A., Müller, M. A., Drosten, C., & Pöhlmann, S. (2020). SARS-CoV-2 Cell Entry Depends on ACE2 and TMPRSS2 and Is Blocked by a Clinically Proven Protease Inhibitor. Cell, 181(2), 271–280.e8.

Hu, B., Huang, S., & Yin, L. (2021). The cytokine storm and COVID-19. Journal of Medical Virology, 93(1), 250–256.

Hussain, S., Pan, J. ‘an Chen, Y., Yang, Y., Xu, J., Peng, Y., Wu, Y., Li, Z., Zhu, Y., Tien, P., & Guo, D. (2005). Identification of novel subgenomic RNAs and noncanonical transcription initiation signals of severe acute respiratory syndrome coronavirus. Journal of Virology, 79(9), 5288–5295.

Islam, A. B. M. M. K., & Khan, M. A.-A.-K. (2020). Lung transcriptome of a COVID-19 patient and systems biology predictions suggest impaired surfactant production which may be druggable by surfactant therapy. Scientific Reports, 10(1), 19395.

Kalfaoglu, B., Almeida-Santos, J., Tye, C. A., Satou, Y., & Ono, M. (2020). T-Cell Hyperactivation and Paralysis in Severe COVID-19 Infection Revealed by Single-Cell Analysis. Frontiers in Immunology, 11, 589380.

Knoblach, B., Ishida, R., Hobman, T. C., & Rachubinski, R. A. (2021). Peroxisomes exhibit compromised structure and matrix protein content in SARS-CoV-2-infected cells. Molecular Biology of the Cell, 32(14), 1273–1282.

Kouhpayeh, H. R., Tabasi, F., Dehvari, M., Naderi, M., Bahari, G., Khalili, T., Clark, C., Ghavami, S., & Taheri, M. (2021). Association between angiotensinogen (AGT), angiotensin-converting enzyme (ACE) and angiotensin-II receptor 1 (AGTR1) polymorphisms and COVID-19 infection in the southeast of Iran: a preliminary case-control study. Translational Medicine Communications, 6(1), 26.

Kuleshov, M. V., Jones, M. R., Rouillard, A. D., Fernandez, N. F., Duan, Q., Wang, Z., Koplev, S., Jenkins, S. L., Jagodnik, K. M., Lachmann, A., McDermott, M. G., Monteiro, C. D., Gundersen, G. W., & Ma’ayan, A. (2016). Enrichr: a comprehensive gene set enrichment analysis web server 2016 update. Nucleic Acids Research, 44(W1), W90–W97.

Lamb, J., Crawford, E. D., Peck, D., Modell, J. W., Blat, I. C., Wrobel, M. J., Lerner, J., Brunet, J.-P., Subramanian, A., Ross, K. N., Reich, M., Hieronymus, H., Wei, G., Armstrong, S. A., Haggarty, S. J., Clemons, P. A., Wei, R., Carr, S. A., Lander, E. S., & Golub, T. R. (2006). The Connectivity Map: using gene-expression signatures to connect small molecules, genes, and disease. Science, 313(5795), 1929–1935.

Langford, B. J., So, M., Raybardhan, S., Leung, V., Westwood, D., MacFadden, D. R., Soucy, J.-P. R., & Daneman, N. (2020). Bacterial co-infection and secondary infection in patients with COVID-19: a living rapid review and meta-analysis. Clinical Microbiology and Infection: The Official Publication of the European Society of Clinical Microbiology and Infectious Diseases, 26(12), 1622–1629.

Langmead, B., & Salzberg, S. L. (2012). Fast gapped-read alignment with Bowtie 2. Nature Methods, 9(4), 357–359.

Lazzaroni, M. G., Piantoni, S., Masneri, S., Garrafa, E., Martini, G., Tincani, A., Andreoli, L., & Franceschini, F. (2021). Coagulation dysfunction in COVID-19: The interplay between inflammation, viral infection and the coagulation system. Blood Reviews, 46, 100745.

Leng, L., Cao, R., Ma, J., Mou, D., Zhu, Y., Li, W., Lv, L., Gao, D., Zhang, S., Gong, F., Zhao, L., Qiu, B., Xiang, H., Hu, Z., Feng, Y., Dai, Y., Zhao, J., Wu, Z., Li, H., & Zhong, W. (2020). Pathological features of COVID-19-associated lung injury: a preliminary proteomics report based on clinical samples. Signal Transduction and Targeted Therapy, 5(1), 240.

Leong, W. F., & Chow, V. T. K. (2006). Transcriptomic and proteomic analyses of rhabdomyosarcoma cells reveal differential cellular gene expression in response to enterovirus 71 infection. Cellular Microbiology, 8(4), 565–580.

Liang, S., Yu, Z., Ma, Y.-Y., Guo, C.-F., Lu, H.-Z., & Xu, J.-F. (2021). Early predicting indicators of conversion from mild to moderate in overseas-imported COVID-19 cases. Annals of Translational Medicine, 9(20), 1584.

Liao, M., Liu, Y., Yuan, J., Wen, Y., Xu, G., Zhao, J., Cheng, L., Li, J., Wang, X., Wang, F., Liu, L., Amit, I., Zhang, S., & Zhang, Z. (2020). Single-cell landscape of bronchoalveolar immune cells in patients with COVID-19. Nature Medicine, 26(6), 842–844.

Liao, Y., Smyth, G. K., & Shi, W. (2014). featureCounts: an efficient general purpose program for assigning sequence reads to genomic features. Bioinformatics, 30(7), 923–930.

Liberzon, A., Subramanian, A., Pinchback, R., Thorvaldsdóttir, H., Tamayo, P., & Mesirov, J. P. (2011). Molecular signatures database (MSigDB) 3.0. Bioinformatics, 27(12), 1739–1740.

Li, H., Handsaker, B., Wysoker, A., Fennell, T., Ruan, J., Homer, N., Marth, G., Abecasis, G., Durbin, R., & 1000 Genome Project Data Processing Subgroup. (2009). The Sequence Alignment/Map format and SAMtools. Bioinformatics, 25(16), 2078–2079.

Liu, N.-N., Ma, Q., Ge, Y., Yi, C.-X., Wei, L.-Q., Tan, J.-C., Chu, Q., Li, J.-Q., Zhang, P., & Wang, H. (2020). Microbiome dysbiosis in lung cancer: from composition to therapy. NPJ Precision Oncology, 4(1), 33.

Liu, Y., O’Brien, J. L., Ajami, N. J., Scheurer, M. E., Amirian, E. S., Armstrong, G., Tsavachidis, S., Thrift, A. P., Jiao, L., Wong, M. C., Smith, D. P., Spitz, M. R., Bondy, M. L., Petrosino, J. F., & Kheradmand, F. (2018). Lung tissue microbial profile in lung cancer is distinct from emphysema. American Journal of Cancer Research, 8(9), 1775–1787.

Love, M. I., Huber, W., & Anders, S. (2014). Moderated estimation of fold change and dispersion for RNA-seq data with DESeq2. Genome Biology, 15(12), 550.

Ma, L., Sahu, S. K., Cano, M., Kuppuswamy, V., Bajwa, J., McPhatter, J., Pine, A., Meizlish, M. L., Goshua, G., Hong Chang, C., Zhang, H., Price, C., Bahel, P., Rinder, H., Lei, T., Day, A., Reynolds, D., Wu, X., Schriefer, R., … Kulkarni, H. S. (2021). Increased complement activation is a distinctive feature of severe SARS-CoV-2 infection. Science Immunology, 6(59). https://doi.org/10.1126/sciimmunol.abh2259

Mason, R. J. (2020). Pathogenesis of COVID-19 from a cell biology perspective. In European Respiratory Journal (Vol. 55, Issue 4, p. 2000607). https://doi.org/10.1183/13993003.00607-2020

Masters, P. S. (2006). The molecular biology of coronaviruses. Advances in Virus Research, 66, 193–292.

McMurdie, P. J., & Holmes, S. (2013). phyloseq: an R package for reproducible interactive analysis and graphics of microbiome census data. PloS One, 8(4), e61217.

Mehrian-Shai, R. (2020). A rational approach to COVID-19. In Human Genomics (Vol. 14, Issue 1). https://doi.org/10.1186/s40246-020-00300-5

Mestriner, F. L. A. C., Spiller, F., Laure, H. J., Souto, F. O., Tavares-Murta, B. M., Rosa, J. C., Basile-Filho, A., Ferreira, S. H., Greene, L. J., & Cunha, F. Q. (2007). Acute-phase protein alpha-1-acid glycoprotein mediates neutrophil migration failure in sepsis by a nitric oxide-dependent mechanism. Proceedings of the National Academy of Sciences of the United States of America, 104(49), 19595–19600.

Millet, J. K., & Whittaker, G. R. (2015). Host cell proteases: Critical determinants of coronavirus tropism and pathogenesis. Virus Research, 202, 120–134.

Moona, A. A., & Islam, M. R. (2021). Mucormycosis or black fungus is a new fright in India during covid-19 pandemic: Associated risk factors and actionable items. In Public Health in Practice (Vol. 2, p. 100153). https://doi.org/10.1016/j.puhip.2021.100153

Mostafa, H. H., Fissel, J. A., Fanelli, B., Bergman, Y., Gniazdowski, V., Dadlani, M., Carroll, K. C., Colwell, R. R., & Simner, P. J. (2020). Metagenomic Next-Generation Sequencing of Nasopharyngeal Specimens Collected from Confirmed and Suspect COVID-19 Patients. mBio, 11(6). https://doi.org/10.1128/mBio.01969-20

Nardacci, R., Colavita, F., Castilletti, C., Lapa, D., Matusali, G., Meschi, S., Del Nonno, F., Colombo, D., Capobianchi, M. R., Zumla, A., Ippolito, G., Piacentini, M., & Falasca, L. (2021). Evidences for lipid involvement in SARS-CoV-2 cytopathogenesis. Cell Death & Disease, 12(3), 263.

Nguyen, T. T., Pathirana, P. N., Nguyen, T., Nguyen, Q. V. H., Bhatti, A., Nguyen, D. C., Nguyen, D. T., Nguyen, N. D., Creighton, D., & Abdelrazek, M. (2021). Genomic mutations and changes in protein secondary structure and solvent accessibility of SARS-CoV-2 (COVID-19 virus). Scientific Reports, 11(1), 3487.

Nienhold, R., Ciani, Y., Koelzer, V. H., Tzankov, A., Haslbauer, J. D., Menter, T., Schwab, N., Henkel, M., Frank, A., Zsikla, V., Willi, N., Kempf, W., Hoyler, T., Barbareschi, M., Moch, H., Tolnay, M., Cathomas, G., Demichelis, F., Junt, T., & Mertz, K. D. (2020). Two distinct immunopathological profiles in autopsy lungs of COVID-19. Nature Communications, 11(1), 5086.

Papoutsoglou, G., Karaglani, M., Lagani, V., Thomson, N., Røe, O. D., Tsamardinos, I., & Chatzaki, E. (2021). Automated machine learning optimizes and accelerates predictive modeling from COVID-19 high throughput datasets. Scientific Reports, 11(1), 15107.

Patel, H., Ashton, N. J., Dobson, R. J. B., Andersson, L.-M., Yilmaz, A., Blennow, K., Gisslen, M., & Zetterberg, H. (2021). Proteomic blood profiling in mild, severe and critical COVID-19 patients. Scientific Reports, 11(1), 6357.

pheatmap: Pretty Heatmaps. (n.d.). Retrieved August 20, 2021, from https://cran.r-project.org/web/packages/pheatmap/index.html

Prakash, H., & Chakrabarti, A. (2021). Epidemiology of Mucormycosis in India. Microorganisms, 9(3). https://doi.org/10.3390/microorganisms9030523

Prakrithi, P., Lakra, P., Sundar, D., Kapoor, M., Mukerji, M., Gupta, I., & The Indian Genome Variation Consortium. (2021). Genetic risk prediction of COVID-19 susceptibility and severity in the Indian population. In bioRxiv. medRxiv. https://doi.org/10.1101/2021.04.13.21255447

Pushkarev, V. V., Sokolova, L. K., Chervyakova, S. A., Belchina, Y. B., Kovzun, O. I., Pushkarev, V. M., & Tronko, M. D. (2021). Plasma Apolipoproteins A1/B and OxLDL Levels in Patients with Covid-19 As Possible Markers of the Disease. Cytology and Genetics, 55(6), 519–523.

Quast, C., Pruesse, E., Yilmaz, P., Gerken, J., Schweer, T., Yarza, P., Peplies, J., & Glöckner, F. O. (2013). The SILVA ribosomal RNA gene database project: improved data processing and web-based tools. Nucleic Acids Research, 41(Database issue), D590–D596.

Rawson, T. M., Moore, L. S. P., Zhu, N., Ranganathan, N., Skolimowska, K., Gilchrist, M., Satta, G., Cooke, G., & Holmes, A. (2020). Bacterial and Fungal Coinfection in Individuals With Coronavirus: A Rapid Review To Support COVID-19 Antimicrobial Prescribing. Clinical Infectious Diseases: An Official Publication of the Infectious Diseases Society of America, 71(9), 2459–2468.

Ripa, M., Galli, L., Poli, A., Oltolini, C., Spagnuolo, V., Mastrangelo, A., Muccini, C., Monti, G., De Luca, G., Landoni, G., Dagna, L., Clementi, M., Rovere Querini, P., Ciceri, F., Tresoldi, M., Lazzarin, A., Zangrillo, A., Scarpellini, P., Castagna, A., & COVID-BioB study group. (2021). Secondary infections in patients hospitalized with COVID-19: incidence and predictive factors. Clinical Microbiology and Infection: The Official Publication of the European Society of Clinical Microbiology and Infectious Diseases, 27(3), 451–457.

Rodriguez, C., de Prost, N., Fourati, S., Lamoureux, C., Gricourt, G., N’debi, M., Canoui-Poitrine, F., Désveaux, I., Picard, O., Demontant, V., Trawinski, E., Lepeule, R., Surgers, L., Vindrios, W., Lelièvre, J.-D., Mongardon, N., Langeron, O., Cohen, J. L., Mekontso-Dessap, A., … Pawlotsky, J.-M. (2021). Viral genomic, metagenomic and human transcriptomic characterization and prediction of the clinical forms of COVID-19. PLoS Pathogens, 17(3), e1009416.

Sadler, A. J., & Williams, B. R. G. (2008). Interferon-inducible antiviral effectors. Nature Reviews. Immunology, 8(7), 559–568.

Saheb Sharif-Askari, N., Saheb Sharif-Askari, F., Mdkhana, B., Al Heialy, S., Alsafar, H. S., Hamoudi, R., Hamid, Q., & Halwani, R. (2021). Enhanced expression of immune checkpoint receptors during SARS-CoV-2 viral infection. Molecular Therapy. Methods & Clinical Development, 20, 109–121.

Sanchez-Cerrillo, I., Landete, P., Aldave, B., Sanchez-Alonso, S., Sanchez-Azofra, A., Marcos-Jimenez, A., Avalos, E., Alcaraz-Serna, A., de Los Santos, I., Mateu-Albero, T., Esparcia, L., Lopez-Sanz, C., Martinez-Fleta, P., Gabrie, L., Del Campo Guerola, L., Calzada, M. J., Gonzalez-Alvaro, I., Alfranca, A., Sanchez-Madrid, F., … Martin-Gayo, E. (2020). Differential Redistribution of Activated Monocyte and Dendritic Cell Subsets to the Lung Associates with Severity of COVID-19. medRxiv : The Preprint Server for Health Sciences. https://doi.org/10.1101/2020.05.13.20100925

Sawicki, S. G., & Sawicki, D. L. (2005). Coronavirus transcription: a perspective. Current Topics in Microbiology and Immunology, 287, 31–55.

Schneider, V. A., Graves-Lindsay, T., Howe, K., Bouk, N., Chen, H.-C., Kitts, P. A., Murphy, T. D., Pruitt, K. D., Thibaud-Nissen, F., Albracht, D., Fulton, R. S., Kremitzki, M., Magrini, V., Markovic, C., McGrath, S., Steinberg, K. M., Auger, K., Chow, W., Collins, J., … Church, D. M. (2017). Evaluation of GRCh38 and de novo haploid genome assemblies demonstrates the enduring quality of the reference assembly. Genome Research, 27(5), 849–864.

Schoggins, J. W., & Rice, C. M. (2011). Interferon-stimulated genes and their antiviral effector functions. Current Opinion in Virology, 1(6), 519–525.

Sen, M., Honavar, S. G., Bansal, R., Sengupta, S., Rao, R., Kim, U., Sharma, M., Sachdev, M., Grover, A. K., Surve, A., Budharapu, A., Ramadhin, A. K., Tripathi, A. K., Gupta, A., Bhargava, A., Sahu, A., Khairnar, A., Kochar, A., Madhavani, A., … members of the Collaborative OPAI-IJO Study on Mucormycosis in COVID-19 (COSMIC) Study Group. (2021). Epidemiology, clinical profile, management, and outcome of COVID-19-associated rhino-orbital-cerebral mucormycosis in 2826 patients in India - Collaborative OPAI-IJO Study on Mucormycosis in COVID-19 (COSMIC), Report 1. Indian Journal of Ophthalmology, 69(7), 1670–1692.

Sexton, N. R., Smith, E. C., Blanc, H., Vignuzzi, M., Peersen, O. B., & Denison, M. R. (2016). Homology-Based Identification of a Mutation in the Coronavirus RNA-Dependent RNA Polymerase That Confers Resistance to Multiple Mutagens. Journal of Virology, 90(16), 7415–7428.

Shu, T., Ning, W., Wu, D., Xu, J., Han, Q., Huang, M., Zou, X., Yang, Q., Yuan, Y., Bie, Y., Pan, S., Mu, J., Han, Y., Yang, X., Zhou, H., Li, R., Ren, Y., Chen, X., Yao, S., … Zhou, X. (2020). Plasma Proteomics Identify Biomarkers and Pathogenesis of COVID-19. Immunity, 53(5), 1108–1122.e5.

Silva, D. L., Lima, C. M., Magalhães, V. C. R., Baltazar, L. M., Peres, N. T. A., Caligiorne, R. B., Moura, A. S., Fereguetti, T., Martins, J. C., Rabelo, L. F., Abrahão, J. S., Lyon, A. C., Johann, S., & Santos, D. A. (2021). Fungal and bacterial coinfections increase mortality of severely ill COVID-19 patients. The Journal of Hospital Infection, 113, 145–154.

Simmons, G., Zmora, P., Gierer, S., Heurich, A., & Pöhlmann, S. (2013). Proteolytic activation of the SARS-coronavirus spike protein: cutting enzymes at the cutting edge of antiviral research. Antiviral Research, 100(3), 605–614.

Singh, R., Hemati, H., Bajpai, M., Yadav, P., Maheshwari, A., Kumar, S., Agrawal, S., Sevak, J. K., Islam, M., Mars, J. S., Sarin, S. K., & Trehanpati, N. (2021). Sustained expression of inflammatory monocytes and activated T cells in COVID-19 patients and recovered convalescent plasma donors. Immunity, Inflammation and Disease, 9(4), 1279–1290.

Smirnova, O. A., Ivanova, O. N., Bartosch, B., Valuev-Elliston, V. T., Mukhtarov, F., Kochetkov, S. N., & Ivanov, A. V. (2016). Hepatitis C Virus NS5A Protein Triggers Oxidative Stress by Inducing NADPH Oxidases 1 and 4 and Cytochrome P450 2E1. Oxidative Medicine and Cellular Longevity, 2016, 8341937.

Sriram, K., & Insel, P. A. (2020). A hypothesis for pathobiology and treatment of COVID-19: The centrality of ACE1/ACE2 imbalance. British Journal of Pharmacology, 177(21), 4825–4844.

Sur, S., Khatun, M., Steele, R., Isbell, T. S., Ray, R., & Ray, R. B. (2021). Exosomes from COVID-19 Patients Carry Tenascin-C and Fibrinogen-β in Triggering Inflammatory Signals in Cells of Distant Organ. International Journal of Molecular Sciences, 22(6). https://doi.org/10.3390/ijms22063184

Tang, Y., Liu, J., Zhang, D., Xu, Z., Ji, J., & Wen, C. (2020). Cytokine Storm in COVID-19: The Current Evidence and Treatment Strategies. Frontiers in Immunology, 11, 1708.

Tay, M. Z., Poh, C. M., Rénia, L., MacAry, P. A., & Ng, L. F. P. (2020). The trinity of COVID-19: immunity, inflammation and intervention. In Nature Reviews Immunology (Vol. 20, Issue 6, pp. 363–374). https://doi.org/10.1038/s41577-020-0311-8

Torchiano, M. (2016). Effsize - a package for efficient effect size computation. https://doi.org/10.5281/zenodo.196082

Travaglini, K. J., Nabhan, A. N., Penland, L., Sinha, R., Gillich, A., Sit, R. V., Chang, S., Conley, S. D., Mori, Y., Seita, J., Berry, G. J., Shrager, J. B., Metzger, R. J., Kuo, C. S., Neff, N., Weissman, I. L., Quake, S. R., & Krasnow, M. A. (2020). A molecular cell atlas of the human lung from single-cell RNA sequencing. Nature, 587(7835), 619–625.

Vasiliou, V., Veselkov, K., Bruford, E., & Reichardt, J. K. V. (2021). Standardized nomenclature and open science in Human Genomics. Human Genomics, 15(1), 13.

Vastrad, B., Vastrad, C., & Tengli, A. (2020). Identification of potential mRNA panels for severe acute respiratory syndrome coronavirus 2 (COVID-19) diagnosis and treatment using microarray dataset and bioinformatics methods. 3 Biotech, 10(10), 422.

Vijay, S., Bansal, N., Rao, B. K., Veeraraghavan, B., Rodrigues, C., Wattal, C., Goyal, J. P., Tadepalli, K., Mathur, P., Venkateswaran, R., Venkatasubramanian, R., Khadanga, S., Bhattacharya, S., Mukherjee, S., Baveja, S., Sistla, S., Panda, S., & Walia, K. (2021). Secondary Infections in Hospitalized COVID-19 Patients: Indian Experience. Infection and Drug Resistance, 14, 1893–1903.

Völlmy, F., van den Toorn, H., Zenezini Chiozzi, R., Zucchetti, O., Papi, A., Volta, C. A., Marracino, L., Vieceli Dalla Sega, F., Fortini, F., Demichev, V., Tober-Lau, P., Campo, G., Contoli, M., Ralser, M., Kurth, F., Spadaro, S., Rizzo, P., & Heck, A., Jr. (2021). A serum proteome signature to predict mortality in severe COVID-19 patients. Life Science Alliance, 4(9). https://doi.org/10.26508/lsa.202101099

Wang, P., Jin, X., Zhou, W., Luo, M., Xu, Z., Xu, C., Li, Y., Ma, K., Cao, H., Huang, Y., Xue, G., Jin, S., Nie, H., & Jiang, Q. (2021). Comprehensive analysis of TCR repertoire in COVID-19 using single cell sequencing. Genomics, 113(2), 456–462.

Wang, S., Yao, X., Ma, S., Ping, Y., Fan, Y., Sun, S., He, Z., Shi, Y., Sun, L., Xiao, S., Song, M., Cai, J., Li, J., Tang, R., Zhao, L., Wang, C., Wang, Q., Zhao, L., Hu, H., … Bian, X.-W. (2021). A single-cell transcriptomic landscape of the lungs of patients with COVID-19. Nature Cell Biology, 23(12), 1314–1328.

Wang, X., Park, J., Susztak, K., Zhang, N. R., & Li, M. (2019). Bulk tissue cell type deconvolution with multi-subject single-cell expression reference. Nature Communications, 10(1), 380.

Wickham, H. (2011). Ggplot2. Springer.

Wood, D. E., Lu, J., & Langmead, B. (2019). Improved metagenomic analysis with Kraken 2. Genome Biology, 20(1), 257.

Wu, F., Zhao, S., Yu, B., Chen, Y.-M., Wang, W., Song, Z.-G., Hu, Y., Tao, Z.-W., Tian, J.-H., Pei, Y.-Y., Yuan, M.-L., Zhang, Y.-L., Dai, F.-H., Liu, Y., Wang, Q.-M., Zheng, J.-J., Xu, L., Holmes, E. C., & Zhang, Y.-Z. (2020). A new coronavirus associated with human respiratory disease in China. Nature, 579(7798), 265–269.

Wu, M., Chen, Y., Xia, H., Wang, C., Tan, C. Y., Cai, X., Liu, Y., Ji, F., Xiong, P., Liu, R., Guan, Y., Duan, Y., Kuang, D., Xu, S., Cai, H., Xia, Q., Yang, D., Wang, M.-W., Chiu, I. M., … Kasper, D. L. (2020). Transcriptional and proteomic insights into the host response in fatal COVID-19 cases. Proceedings of the National Academy of Sciences of the United States of America, 117(45), 28336–28343.

Xiong, Y., Liu, Y., Cao, L., Wang, D., Guo, M., Jiang, A., Guo, D., Hu, W., Yang, J., Tang, Z., Wu, H., Lin, Y., Zhang, M., Zhang, Q., Shi, M., Liu, Y., Zhou, Y., Lan, K., & Chen, Y. (2020). Transcriptomic characteristics of bronchoalveolar lavage fluid and peripheral blood mononuclear cells in COVID-19 patients. Emerging Microbes & Infections, 9(1), 761–770.

Yu, G., Wang, L.-G., Han, Y., & He, Q.-Y. (2012). clusterProfiler: an R package for comparing biological themes among gene clusters. Omics: A Journal of Integrative Biology, 16(5), 284–287.

Zhang, H., Ai, J.-W., Yang, W., Zhou, X., He, F., Xie, S., Zeng, W., Li, Y., Yu, Y., Gou, X., Li, Y., Wang, X., Su, H., Xu, T., & Zhang, W. (2020). Metatranscriptomic Characterization of COVID-19 Identified A Host Transcriptional Classifier Associated With Immune Signaling. Clinical Infectious Diseases: An Official Publication of the Infectious Diseases Society of America. https://doi.org/10.1093/cid/ciaa663

Zhou, Z., Ren, L., Zhang, L., Zhong, J., Xiao, Y., Jia, Z., Guo, L., Yang, J., Wang, C., Jiang, S., Yang, D., Zhang, G., Li, H., Chen, F., Xu, Y., Chen, M., Gao, Z., Yang, J., Dong, J., … Wang, J. (2020). Heightened Innate Immune Responses in the Respiratory Tract of COVID-19 Patients. Cell Host & Microbe, 27(6), 883–890.e2.

Zolfaghari Emameh, R., Nosrati, H., Eftekhari, M., Falak, R., & Khoshmirsafa, M. (2020). Expansion of Single Cell Transcriptomics Data of SARS-CoV Infection in Human Bronchial Epithelial Cells to COVID-19. Biological Procedures Online, 22, 16.

