## Supplementary File 1 for "Molecular signature of postmortem lung tissue from COVID-19 patients suggests distinct trajectories driving mortality"

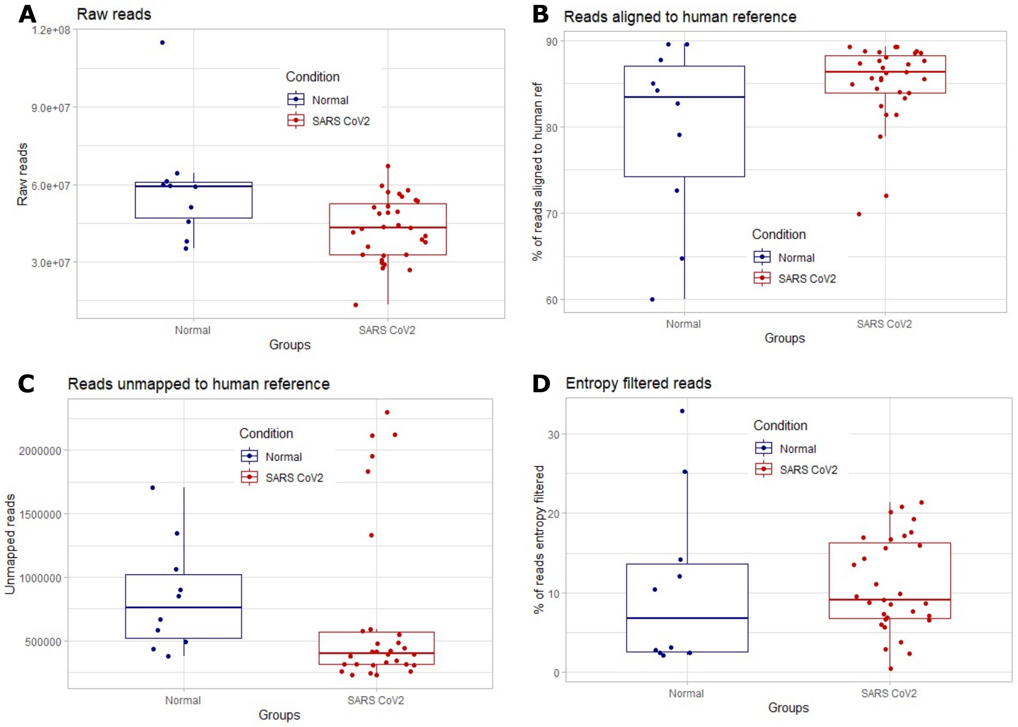


Figure S1 **Distribution of reads in various samples at various steps of the host transcriptome and metatranscriptome analysis**.
**A.** Raw reads distribution before quality control filter **B.** Distribution of reads aligned to human reference after quality control filtering using star aligner **C.** Distribution of number of reads not mapped to human reference after quality control filtering **D.** distribution of percentage of unmapped reads lost after entropy filtering

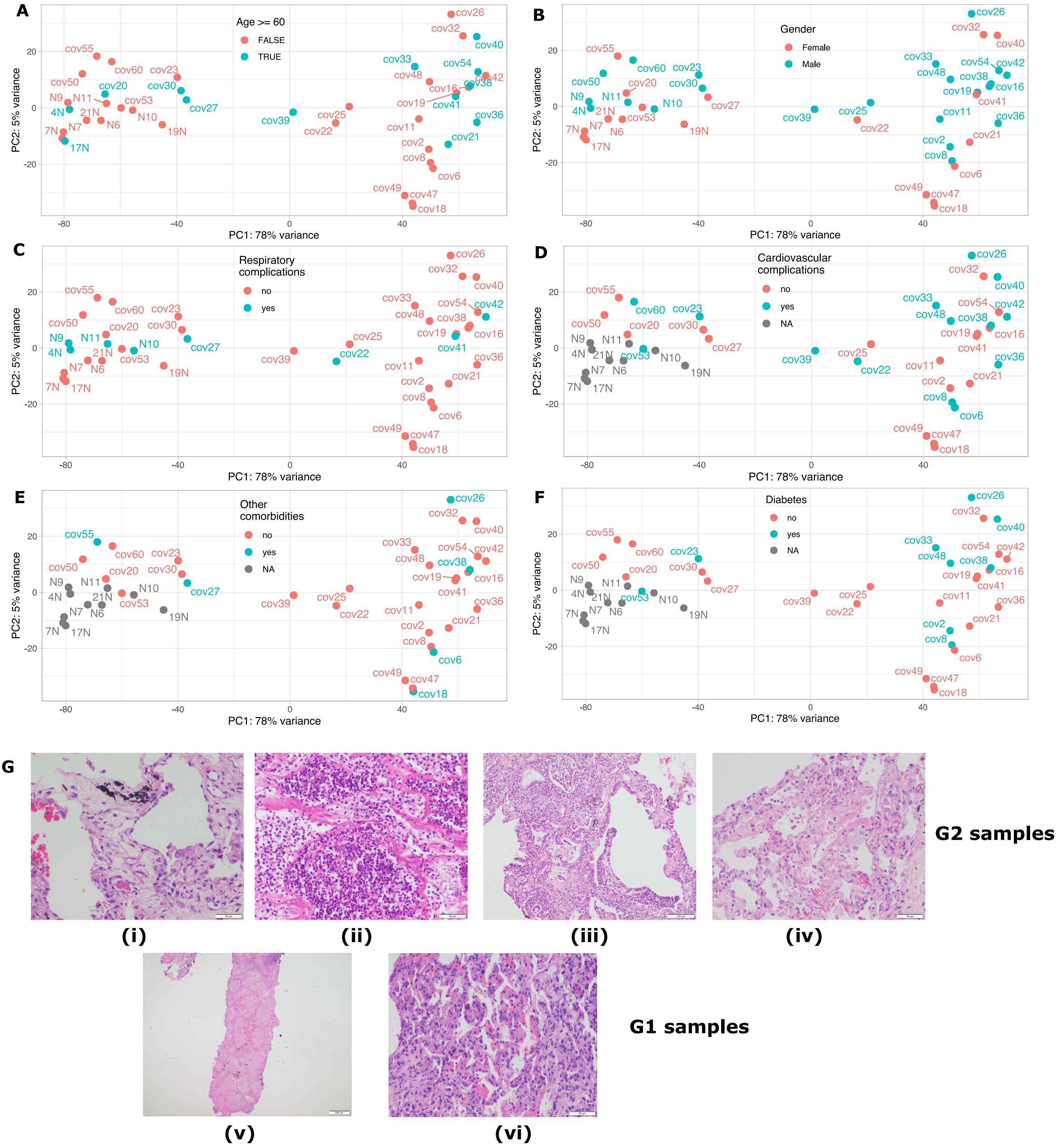


Figure S2 **PCA plots with multiple covariates, Hematoxylin and Eosin stained COVID samples**

**A.** Samples segregated with respect to age. **B.** samples segregated with respect to diabetes. **C.** Samples segregated with respect to liver or kidney complications **D.** Samples segregated with respect to cardio-vascular complication. **E.** Samples segregated with respect to other comorbidities. **F.** Samples segregated with respect to diabetes. **G.** Representative histological images of postmortem lung tissue sections for two patient groups. (i) Photomicrograph shows dilated alveolus with prominence of type II pneumocytes and fibrin in one of the adjoining small blood vessels x200. (ii) Alveolar spaces are filled with neutrophiic exudate and forming microabscesses x200. (iii) Photomicrograph shows dilated small airways and alveolar spaces, filled with mucous and neutrophiic exudate (acute bronchopneumonia) x100. (iv) Photomicrograph shows diffuse alveolar damage with hyaline membrane x200. (v) Low magnification of lung parenchyma in which alveolar spaces are filled with necrotic exudate with bacterial colonies x40. (vi). Photomicrograph shows reactive type II pneumocytes lining alveolar spaces with congested inter-alveolar capillaries x400

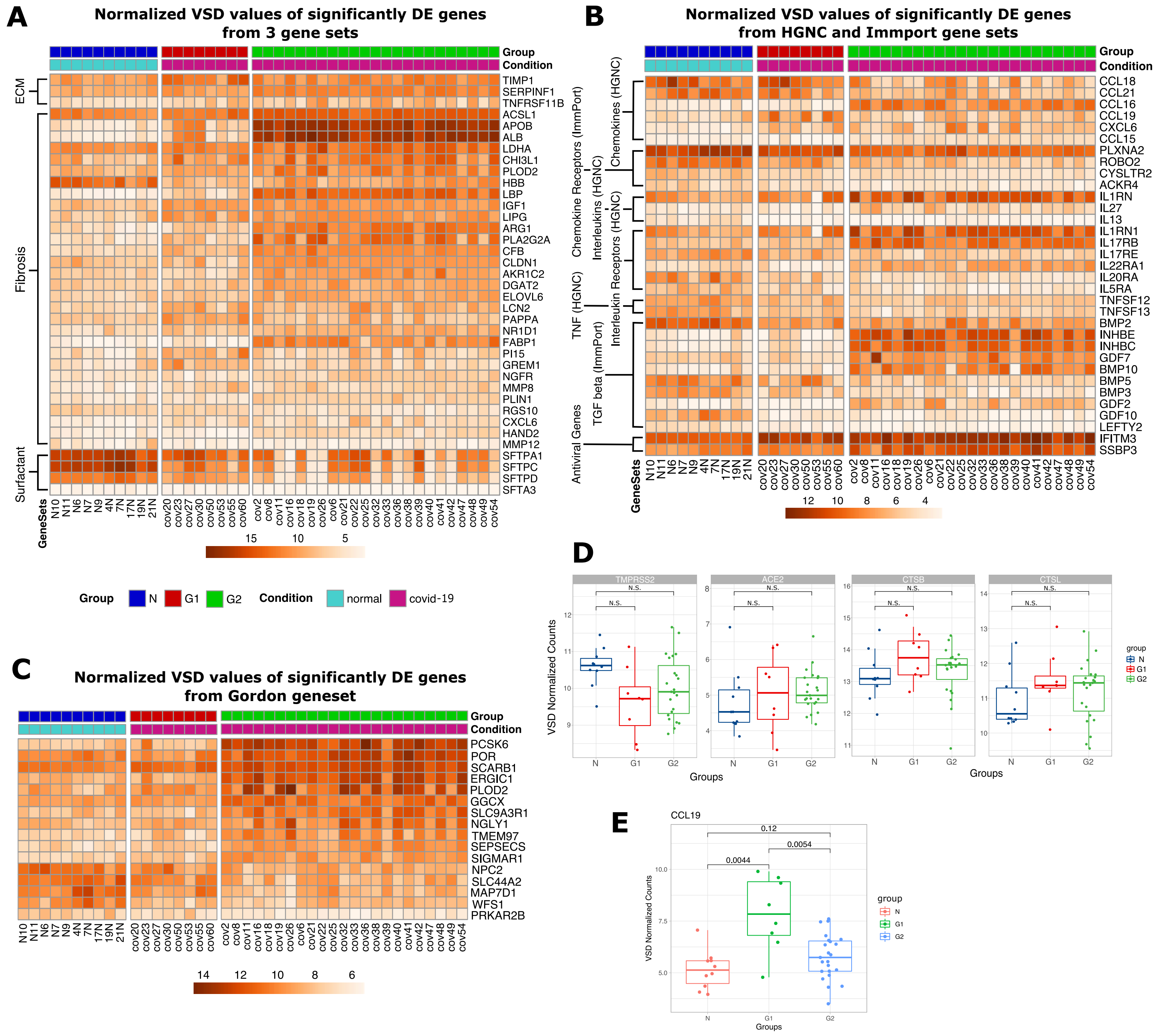


Figure S3

**A.** Heatmap showing normalized expression values for significantly differentially expressed genes from gene sets containing extracellular matrix (ECM) proteins, proteins involved in fibrosis and surfactant proteins. **B.**Heatmap showing normalized expression values for significantly differentially expressed genes from some of the above mentioned pathways. **C.**Heatmap showing normalized expression values for significantly differentially expressed genes from a gene set of host cell proteins found to interact with SARS-CoV-2. **D.** Expression distribution plots of multiple genes in G1 vs G2 vs N, which are essential in the corona virus survival and transmission. **E.** Expression distribution plot of CCL19 which serve as a COVID-19 biomarker

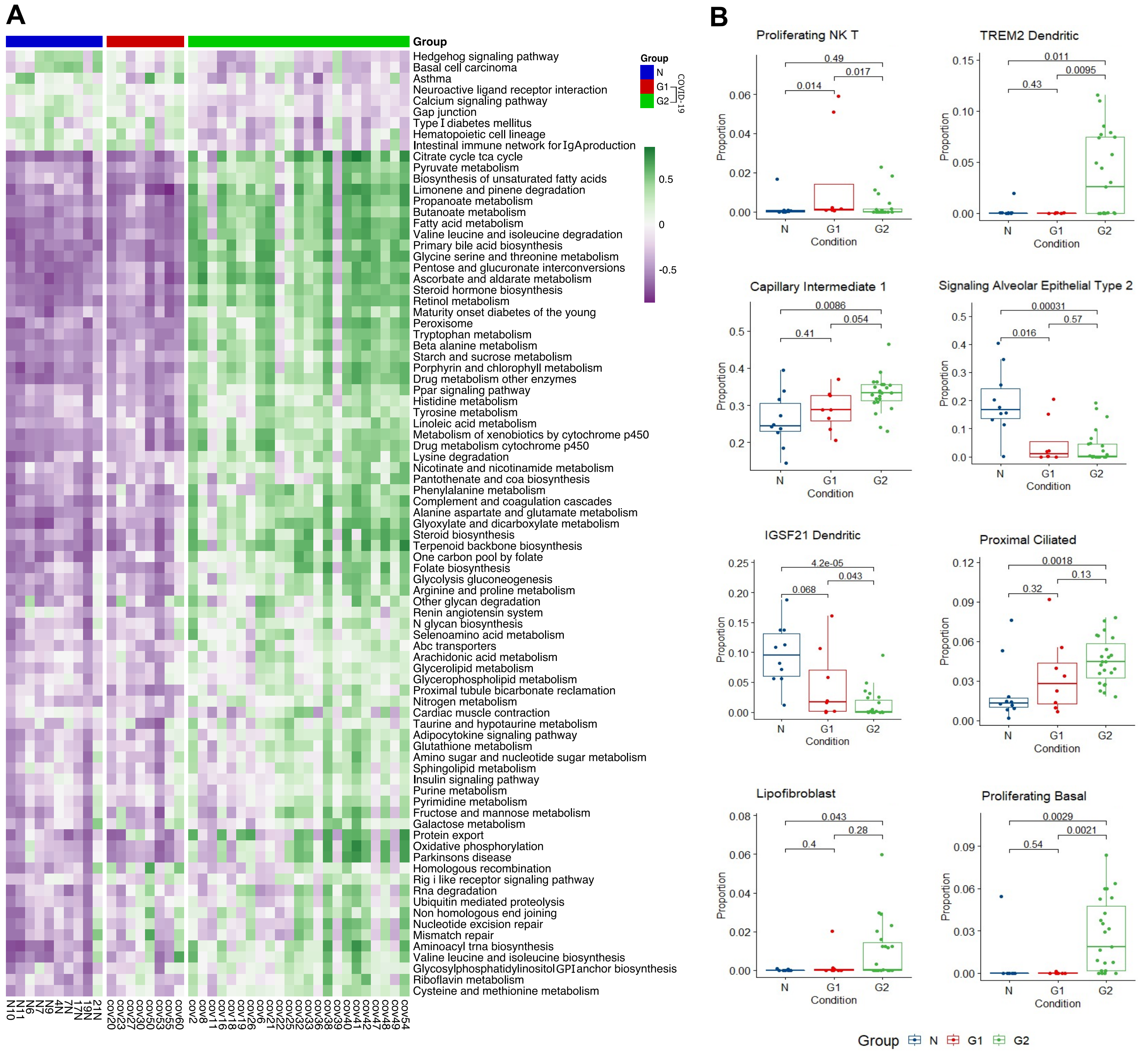


Figure S4 **Gene Set Variation Analysis (GSVA) and cell type deconvolution analysis**

**A.** Heatmap of significantly differentially regulated pathways.produced using KEGG subset of C2 canonical pathways (CP) dataset from MSigDB **B.** Boxplots of significantly relatively altered cell types in comparison to N vs G1 vs G2 samples, produced from results of cell type deconvolution analysis

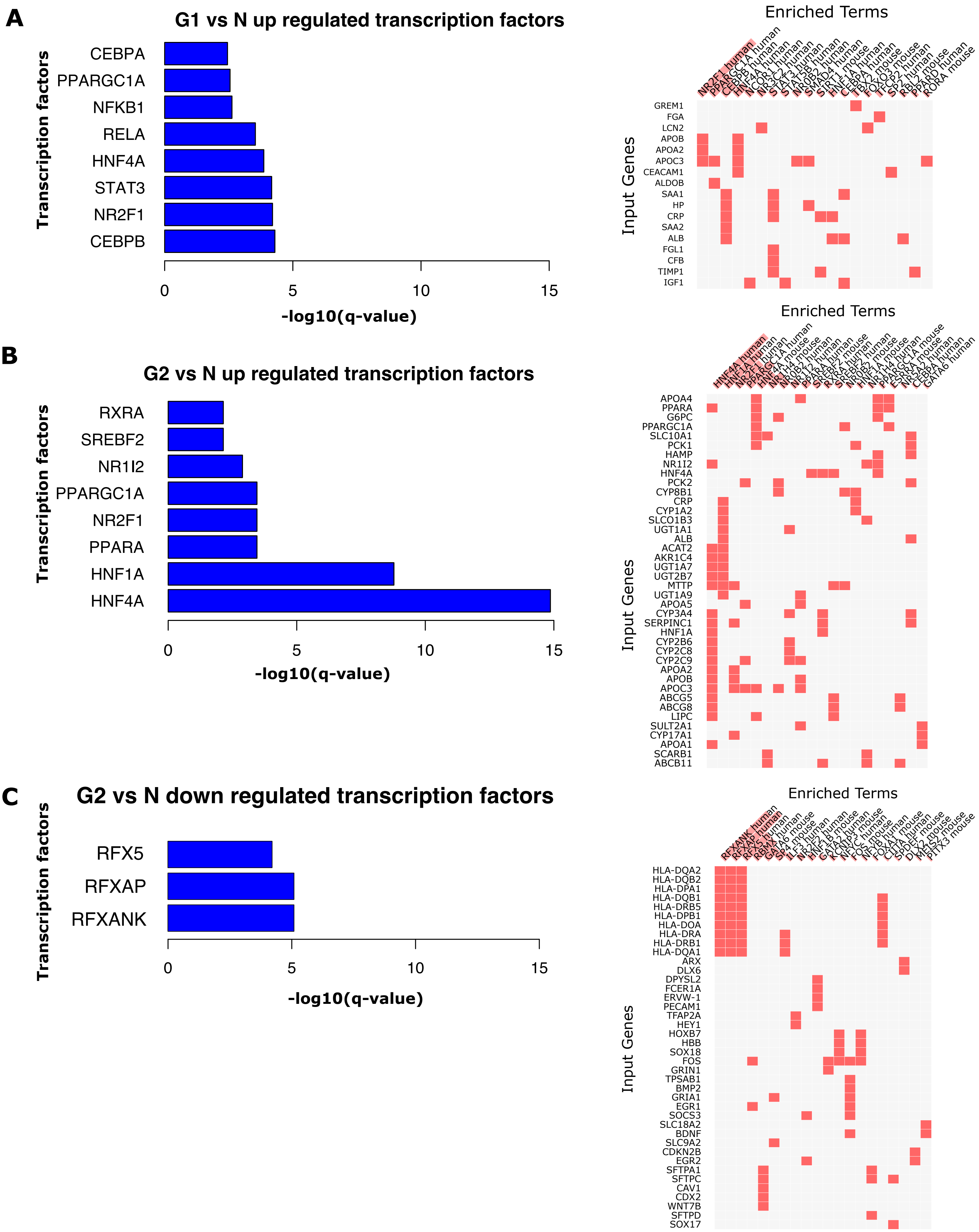


Figure S5. **Summary of transcription factor (TF) analysis**

Barplot shows the transcription factors that regulate most of the significantly differentially expressed genes. The clustergram contains enriched transcription factors on x-axis and its target genes which are differentially regulated on Y-axis **A.** Transcription factors of G1 vs N up regulated genes. **B.** Transcription factors of G2 vs N up regulated genes. **C.** Transcription factors of G2 vs N down regulated genes.

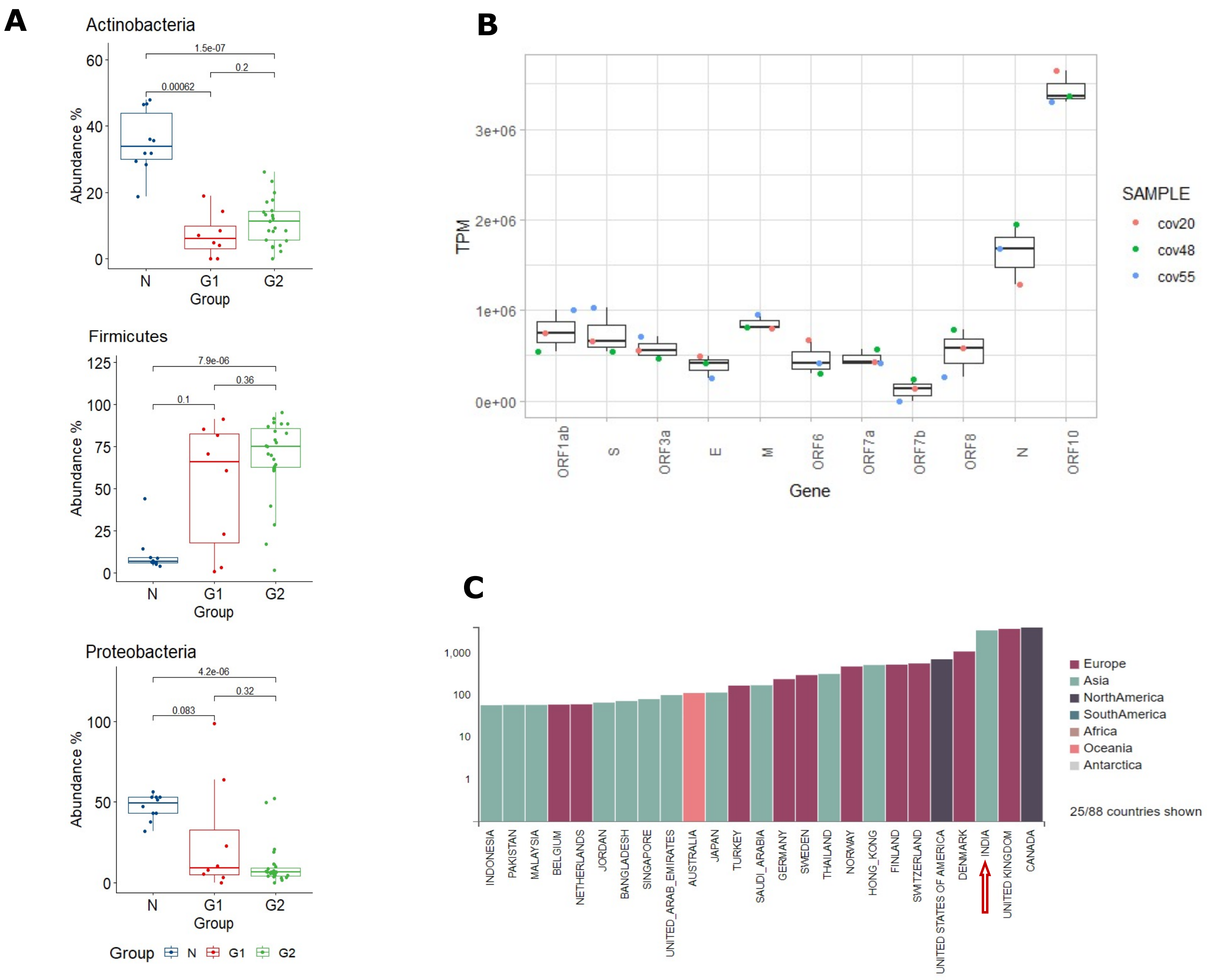


Figure S6 **Metatranscriptome analysis**

**A.** Distribution of a few significantly differentially abundant bacterial phyla in COVID-19 and control samples

**B.** Expression profile of multiple genes in corona virus from three complete genome sequenced corona virus samples.

**C.** Number of viral genomes per country in PANGO database which fall into lineage B.1.36.

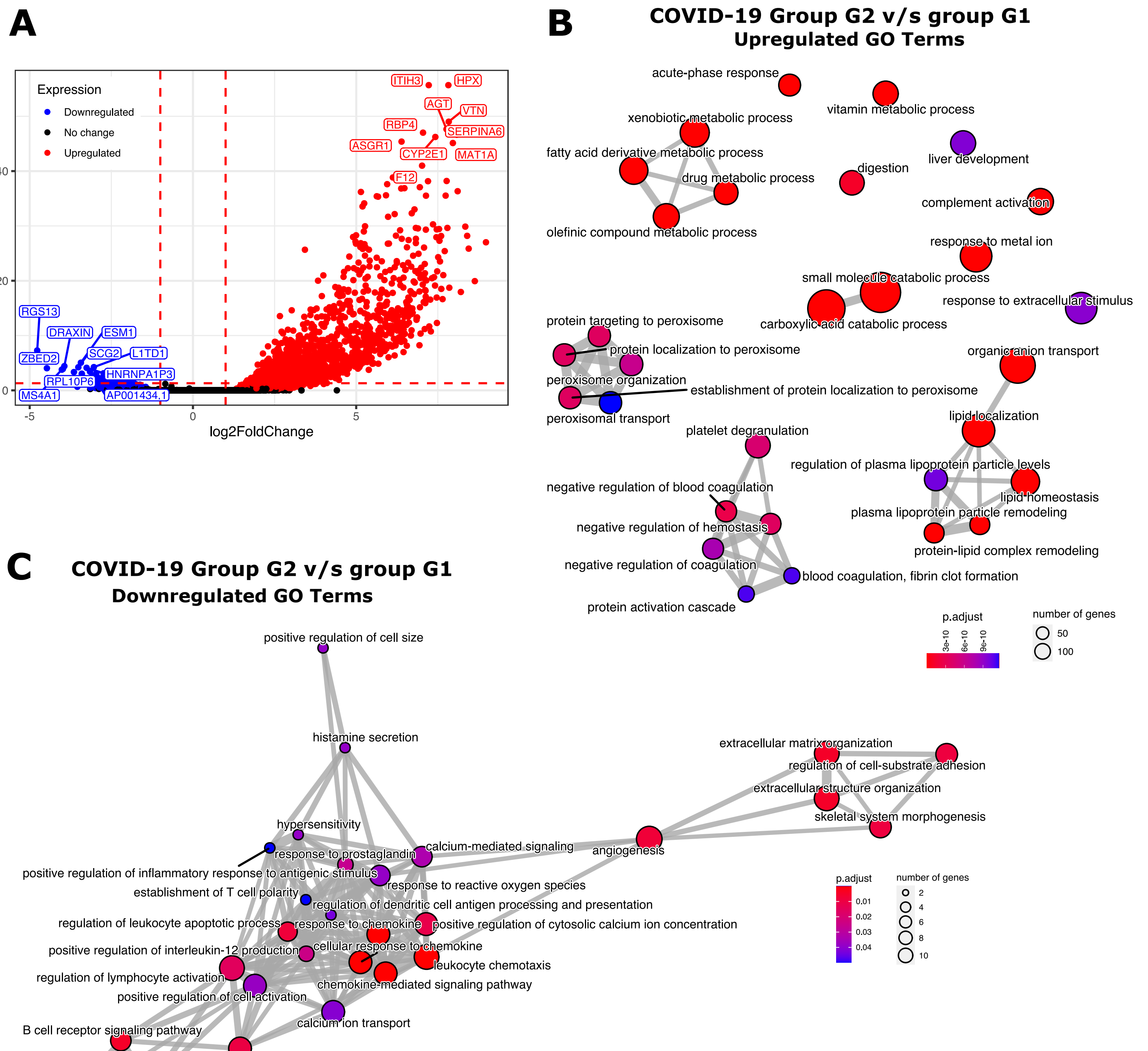


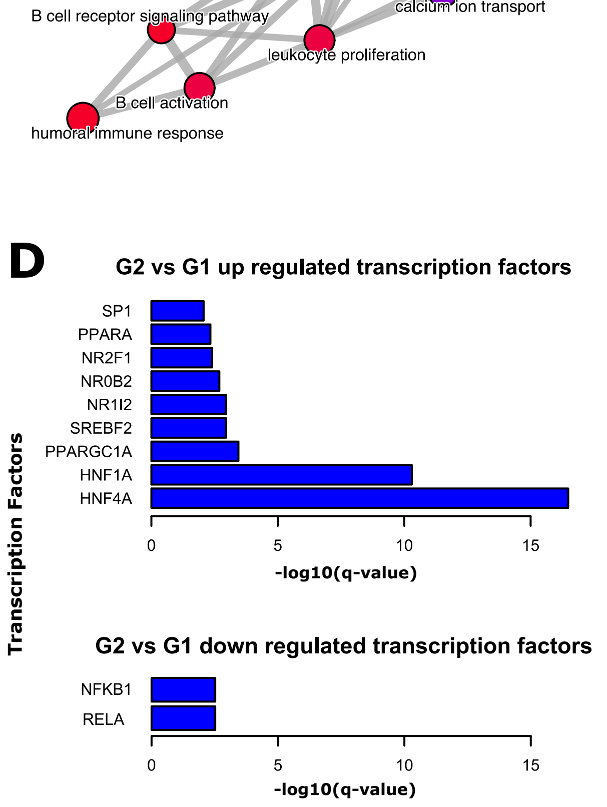


Figure S7 **Significantly differentially expressed genes, GO enrichment and Transcription Factor analysis for G2 v/s G1 comparison**

**A.** Volcano Plot depicting the top 10 significantly upregulated (red) and downregulated (blue) genes when comparing G2 with G1 group of COVID-19 samples using their Fold change (along x-axis) and FDR adjusted p-values (along y-axis).
**B.** GO enrichment maps where the nodes represent enriched biological processes and the edges represent overlapping gene sets for upregulated genes of G2 v/s G1 comparison. As observed, upregulated pathways in G2, compared to G1 include drug metabolic processes along with lipid metabolism and homeostasis along with negative regulation of coagulation and peroxisomal transport. **C.** GO enrichment maps where the nodes represent enriched biological processes and the edges represent overlapping gene sets for downregulated genes of G2 v/s G1 comparison. The pathways that are downregulated in G2 (but relatively upregulated or enriched for G1 samples) are related to humoral immune response, chemokine signaling pathways, angiogenesis and extracellular structure organization. **D.** Barplots indicating the transcription factor targets enriched in significantly up and downregulated genes in G2 vs G1 comparison.
